# Evaluating Aggregated Gene Level eQTL Scores

**DOI:** 10.64898/2026.08.21.746287

**Authors:** Douglas Meyer, Noa Popko, David Laub, Phil Schofield, Tiffany Amariuta, Ludmil B. Alexandrov, Hannah Carter

## Abstract

Genetic feature engineering, used in methods such as transcriptome-wide association study, supports gene-trait association testing by aggregating single variants into gene-level features predictive of expression. To evaluate how different model architectures, LD filtering thresholds, and variant prioritization methods affect expression prediction quality, we trained over 3 million models and evaluated their performance in independent cohorts. Using the best performing models to impute expression and immunotherapy response as an example trait, we found a significant association with the reactive oxygen species pathway (p=0.032). Our model training workflow will support genetic feature engineering towards improved complex trait modeling.

## Background

Genetic modulation of gene expression is commonly tissue^1–3^ and cell-type^4–8^ specific. The identification of variants that modulate gene expression, or expression quantitative trait loci (eQTLs), has supported putative causal gene discovery in many disease contexts such as asthma, diabetes and cancer^9–12^. When used in conjunction with eQTL resources, large cohorts with both genetic and phenotypic measurements – such as UK Biobank^13^ and All of Us^14^ – can nominate genes contributing to a trait of interest^15,16^. Recent efforts to more comprehensively identify eQTLs, like eQTLGen^17^, have largely focused on blood and blood-derived cells. As a small fraction of blood-derived eQTLs generalize to other tissues^17^, the Genotype-Tissue Expression (GTEx) project, which provides eQTLs across multiple different tissue contexts^18,19^ remains useful.

A major challenge to genetic modeling is the large number of previously identified tissue and cell-type-specific eQTL. This has motivated the development of frameworks to aggregate eQTL effects together into gene-level features. PrediXcan^20^ established one of the first approaches for integrating eQTL information by first fitting an elastic net eQTL-based model to impute gene expression towards transcriptome-wide association study (TWAS). Another early TWAS method, FUSION^21^, uses the best performing model of Bayesian sparse linear mixed models (BSLMM), best linear unbiased prediction (BLUP), top predictive eQTL (top1), LASSO, and elastic net.

Subsequent efforts to improve gene-trait association testing have examined alternative model architectures. Using a principal component (PC) regression approach developed for MAGMA^22^, e-MAGMA reported superior causal gene identification to PrediXcan and FUSION^23^. Despite these advances, model performance has been reported to vary substantially across studies and specific tissue contexts. For instance, Grinberg and Wallace^24^ found random forest (RF) models outperformed alternatives in predicting B cell and monocyte expression while Okoro *et al.* found that elastic net generally outperformed other tested model types, including RF, in predicting monocyte gene expression^25^. While these alternative model architectures provide some benefit in specific contexts, benchmarking their performance in independent cohorts remains necessary to evaluate their utility.

Feature selection prior to gene-level aggregation can also improve the expression imputation quality. Finemapping approaches like sum of single effects (SuSiE^26^) or deterministic approximation of posteriors (DAP^27^) identify likely causal variants, significantly improving expression prediction performance^28^. Sequence to function models such as AlphaGenome^29^ can also nominate likely causal variants, though their impact on expression prediction has not been fully characterized. Finally, most existing expression modeling approaches assume regularization (FUSION and PrediXcan) or principal component analysis (MAGMA) adequately handles eQTLs in linkage disequilibrium (LD). Simple approaches to LD filtering, such as PLINK2^30^ LD pruning, may improve out-of-cohort model performance by reducing multicollinearity and model complexity. However, this must be evaluated to justify inclusion of LD filtering for expression modeling.

We developed the EAGLES pipeline to facilitate systematic training and evaluation of gene-level eQTL scoring methods, allowing us to better evaluate the consequences of different model architectures, causal variant nomination strategies, and LD filtering thresholds on expression prediction. Models were trained across multiple GTEx tissue types and benchmarked by their performance in independent cohorts. Whole blood models were validated using the Geuvadis cohort^31^, and other tissue models were validated with normal samples from the Cancer Genome Atlas (TCGA^32^).

## Results

### EAGLES overview and study design

In this study, design choices for constructing gene-level polygenic eQTL scores were evaluated through the newly developed **E**valuation of **A**ggregated **G**ene-**L**evel **e**QTL **S**cores (EAGLES) pipeline. We trained models to predict gene expression from collections of expression-associated SNPs, evaluating model performance sensitivity to 3 main parameters: (i) model architecture, (ii) varying linkage disequilibrium (LD) filtering thresholds, and (iii) different eQTL detection and prioritization methods. An overview of EAGLES is presented in **Fig. 1A**. For each set of parameters, we trained models using GTEx individuals and evaluated their performance in a held-out subset of the GTEx cohort or in an independent, tissue-matched validation cohort (**Table S1**). In total we trained 3,180,897 models covering 23,686 genes across 6 GTEx tissue types (**Fig. 1B, Table S2**). For each gene, we evaluated 4 model types: elastic net (EN), a simple sum of affect allele dosage we hereby call FlipAllele (FA), principal component regression (PCR) and XGBoost (XGB). We considered 4 LD options: no filtering, 100 kb window and 0.8 threshold, 200 kb window and 0.5 threshold, and 500 kb window and 0.2 threshold. We labeled these “LDNone”, “LDLax”, “LDMed”, and “LDStrict”, respectively. Finally, we considered 4 options for defining eQTL per tissue: v8 downloadable eQTLs from GTEx (eQTL), v10 downloadable SuSiE finemapped eQTLs from GTEx (FM), a functional subset of GTEx eQTLs, prioritized by AlphaGenome (AG), and the union of AlphaGenome and SuSiE eQTL tables (AG_or_FM).

**Figure 1:**
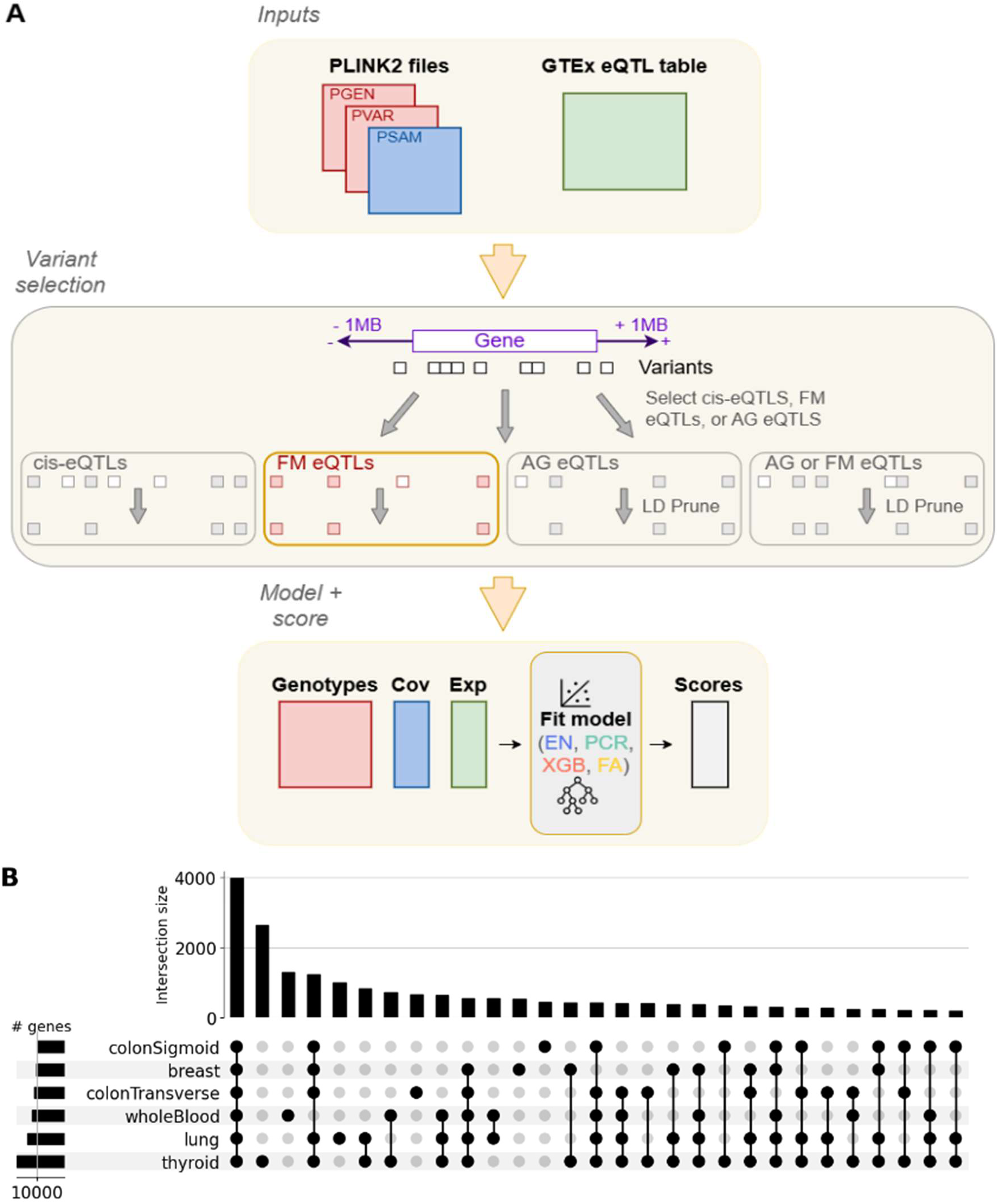
Overview of EAGLES. (a) Overview of the EAGLES pipeline. Genotype data and GTEx eQTL tables serve as the inputs. Variants within ±1 Mb of each gene are selected by the user to be cis-eQTLs, fine-mapped eQTLs, or AlphaGenome-prioritized eQTLs with optional levels of LD pruning. Genotypes, covariates, and expression values are then used to train the user’s chosen model type, producing gene-level scores as the output. (b) UpSet plot summarizing the number of genes per tissue for which eQTLs were available. FM=finemapped, AG=AlphaGenome, EN=elastic net, PCR=principal component regression, XGB=xgboost, FA=flip allele count, LD=linkage disequilibrium; Cov=covariates; Exp=expression.

### Selecting best EAGLES parameters

We started by curating or generating four sets of eQTLs for each tissue type. We downloaded v8 eQTLs (“eQTL”), and v10 SuSiE finemapped eQTLs (“FM”) from GTEx. Next, we utilized AlphaGenome to prioritize eQTLs with strong functional evidence across RNA-seq, DNase-seq, and CAGE modalities to obtain a refined set of germline variants. Finally, we combined the AG and FM tables to include all eQTLs prioritized by finemapping or our AlphaGenome filter (“AG_or_FM”). For each studied tissue and eQTL table, we used held-out GTEx samples to evaluate the consequence of model architecture and LD threshold parameters on EAGLES model performance, using adjusted R^2^ to account for varying numbers of features between EAGLES models. Due to high collinearity among GTEx eQTLs, we did not analyze the “None” LD threshold for that eQTL type. Similarly, due to the reduced number of eQTLs per gene relative to other eQTL sources, we did not assess LDMed or LDStrict thresholds for the FM eQTL type.

For blood, we found that the EN-LDStrict EAGLES mode maximized performance for the most genes (16.3%), representing an incremental increase over other EN or LDStrict EAGLES modes (**Fig 2A**). We considered the possibility that our AG filter preferentially selected an LD independent subset of GTEx eQTLs, so we included “None” LD for evaluation of EAGLES AG models. We found that the EN-LDLax mode maximized performance for the most genes in blood (15%), suggesting that the AG filter reduces but does not eliminate the benefit of LD pruning on model performance (**Fig 2B**). As we expected SuSiE finemapping to converge on LD independent variants, we only examined “None” and “Lax” LD for EAGLES FM models. For blood, we found that EN-LDNone mode maximized performance for the most genes (16.7%), but a notably higher proportion of genes were modeled by a single variant (>71%) than for other eQTL sources (**Fig 2C**). Finally, for EAGLES AG_or_FM models we found that EN-LDMed maximized performance for the most genes (13.9%) with EN-LDLax nearly tied (**Fig 2D**). Overall, we saw that no model configuration was universally optimal among all genes, and that the eQTL source informed the best mode.

**Figure 2:**
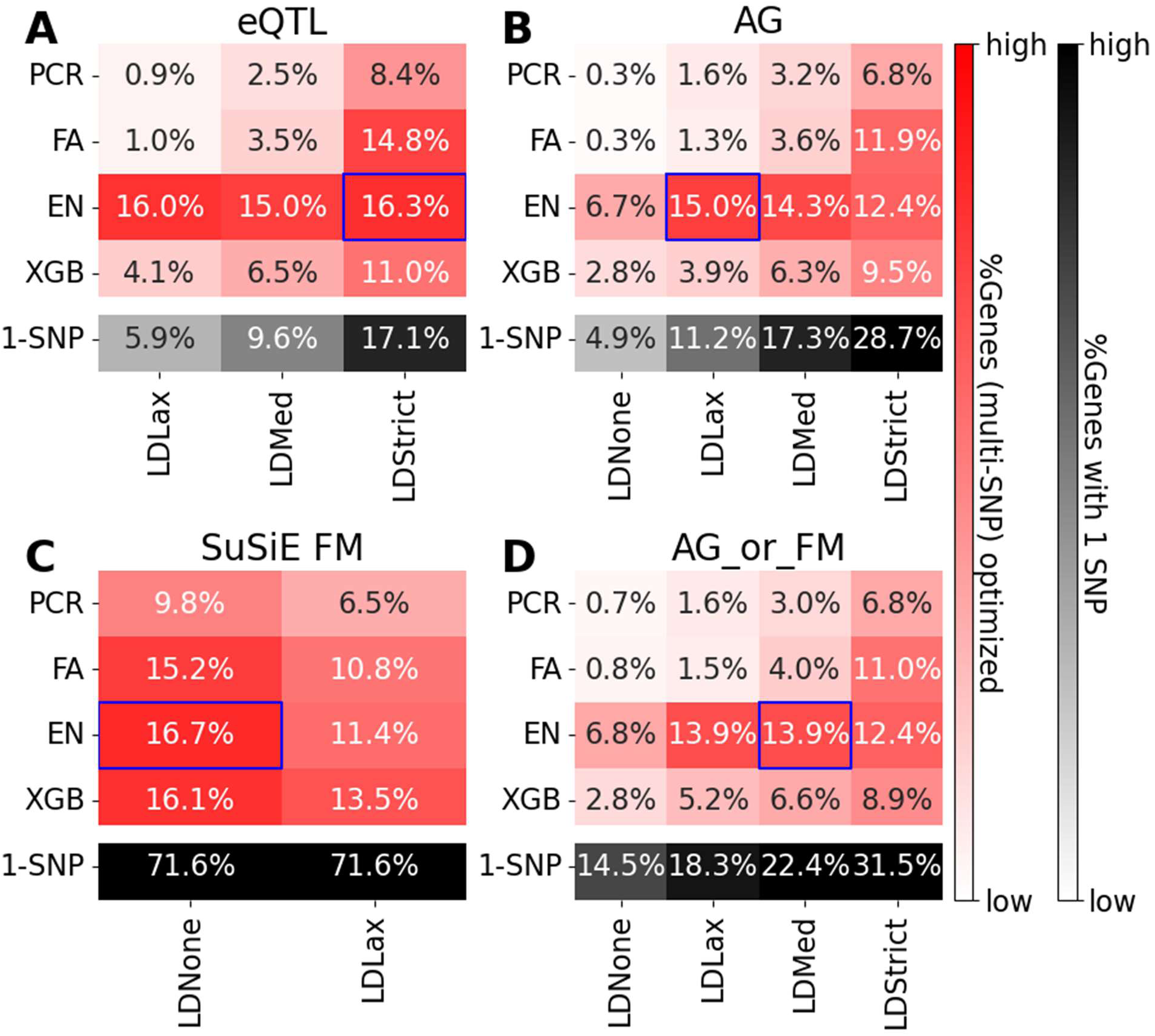
Summary of EAGLES modes impact on adjusted R^2^ in held-out GTEx blood samples. EAGLES models were trained using GTEx eQTLs (A), AlphaGenome filtered eQTLs (B), SuSiE finemapped eQTLs (C) or the union of AlphaGenome and finemapped eQTLs (D). Heatmaps show the fraction of polygenic EAGLES models which maximize performance (adjusted R^2^) relative to other tested EAGLES modes. The black row indicates the fraction of models with a single SNP. Blue boxes indicate the best EAGLES mode. PCR=principal component regression, FA=flip allele count, EN=elastic net, XGB=xgboost, LD=linkage disequilibrium, FM=finemapped, AG=AlphaGenome.

Consistent with our findings in blood, no single EAGLES mode emerged as uniformly dominant in any of the examined tissue types. The best EAGLES mode optimizes performance in at most 16.5% (eQTL), 15.7% (AG), 18.2% (FM) and 14.4% (AG_or_FM) of genes (**Fig S1-5**). From this, we considered determination of the best EAGLES mode on a gene-by-gene basis. However, this yielded minor performance improvements (**Fig S6**).

As a result, we decided to proceed in our analysis using the single best EAGLES mode for each tissue and eQTL type pair. While EN was the most commonly chosen model type (21/24 pairs), XGB (2/24) was also chosen, suggesting nonlinear architectures like XGB may be useful for expression imputation in specific contexts (**Fig S7**).

From the analysis of performance in held-out samples, we consistently found the best performing EAGLES AG and AG_or_FM models involved some LD pruning, while the best performing FM models involved no LD pruning. We next compared performance between models with no LD pruning and models with LD pruning in the independent cohorts to better quantify the impact of LD pruning on EAGLES model performance. Consistent with our findings in held-out samples, we found significant improvements in model performance resulting from LD pruning across all examined tissues (**Fig S8**).

### Identification of high-performance EAGLES genes

Cis-heritability (h^2^) of gene expression is a frequently used metric to assess the performance potential (R^2^) of expression prediction models^20,21^, setting an upper bound on the variance in expression attributable to eQTLs. By evaluating the trade-off between model performance and model complexity, and identifying potentially overfit models, the F-test is another useful metric to evaluate regression model performance^33^. We used the GTEx cohort to stratify genes into four groups based on presence of significant cis-heritable expression (GCTA-h^2^ p<0.05) and outcome of the F-test. As expected, we consistently observed highest performance (adjusted R^2^) in the independent blood cohort (Geuvadis) for genes satisfying both criteria, and low performance in genes satisfying neither criteria (**Fig 3**). We observed similar trends in the other examined tissues (**Fig S9-S13**).

**Figure 3:**
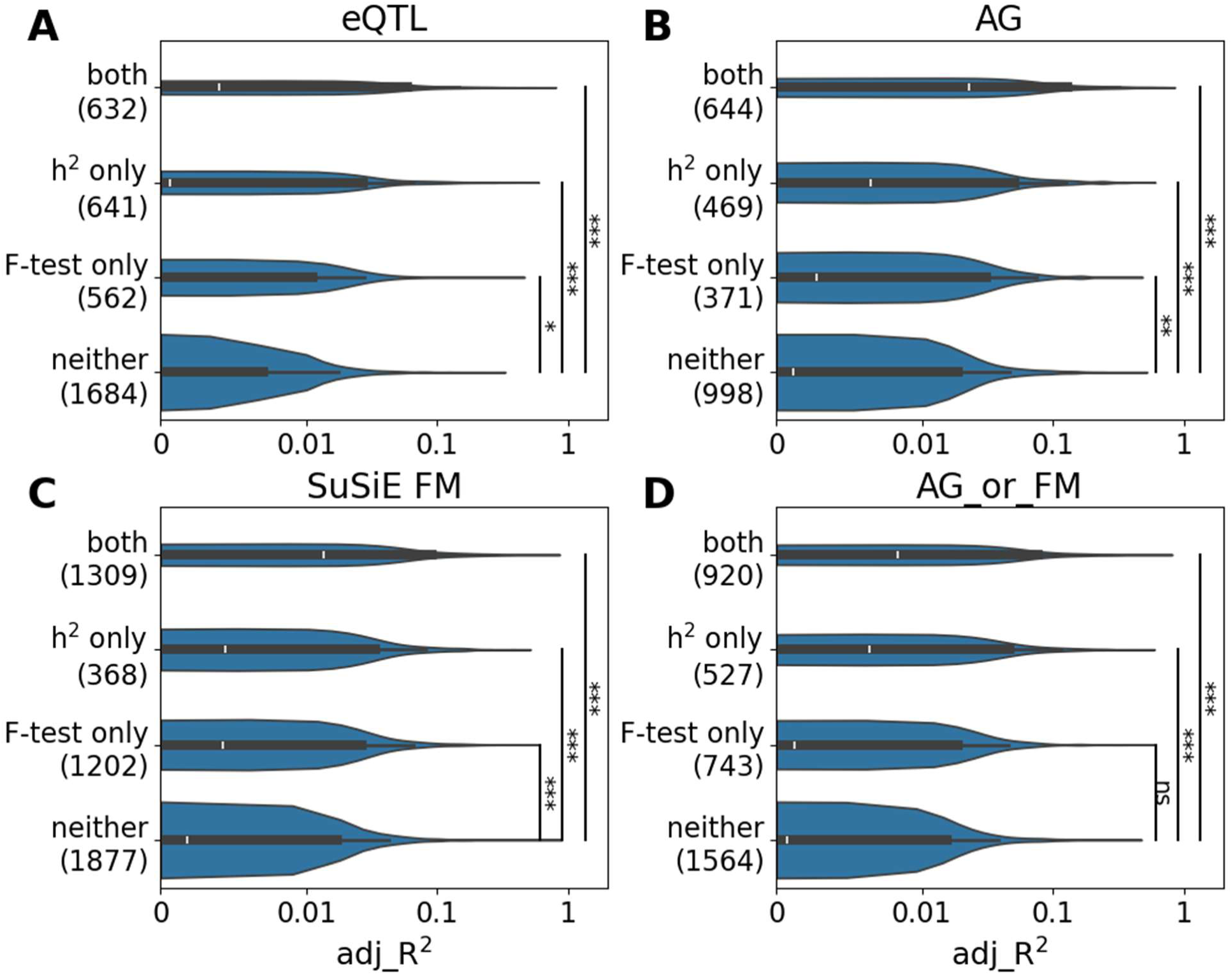
Identifying high-performing EAGLES models in the independent blood cohort. EAGLES models use features from GTEx eQTLs (A), AlphaGenome prioritized eQTLs (B), GTEx finemapped (FM) eQTLs (C) or the union of AlphaGenome and finemapped (AG_or_FM) eQTLs (D). Models were grouped based on whether they passed the F test and/or predicted expression of genes with significant cis-heritability estimated in GTEx. Labels on y-axis note F-test and heritability (h^2^) category with the number of genes per category in parenthesis. Mann-Whitney-U tests (left-tailed) were used for comparisons: ***<0.001; **<0.01; *<0.05; ns>0.05. AG=AlphaGenome; FM=finemapped; h^2^=significant cis-heritability

As expected, EAGLES performance was poorest in genes with no significant heritable expression. However, we observed high performance (adjusted R^2^ > 0.2) in 229 models. To assess if this could be explained by heritability differences across cohorts, we estimated h^2^ in Geuvadis and GTEx to split genes into 4 groups: h^2^>0 in neither, in GTEx only, in Geuvadis only, or in both cohorts. When comparing EAGLES performance in Geuvadis, we saw better performance in genes with significant h^2^ in Geuvadis than genes with significant h^2^ in GTEx only (**Fig S14**). Of the 229 high performing genes that showed no significant cis heritability in GTEx, 226 had significant measurable h^2^ in Geuvadis. Because of these differences in heritability between cohorts, estimated h^2^ may only be a useful predictor of EAGLES performance in genes with significant heritability in both cohorts.

### eQTL source and EAGLES performance in high-performance genes

As h^2^ has been reported as an upper bound on model performance^20,21^, we considered the relationship between h^2^ and adjusted R^2^ for EAGLES models using different types of eQTLs. We focused this analysis on the 3 high performing groups of genes identified earlier (passing F-test, h^2^>0, or both). Gene groups were selected based on metrics computed in GTEx and we quantified the relationship between h^2^ and adjusted R^2^ in the independent cohort. By permutation test, we found a significantly stronger relationship between h^2^ and adjusted R^2^ for the F-test-only genes in AG EAGLES than in SuSiE FM EAGLES (**Fig 4A**). For genes with significant heritability, we saw no significant differences in the h^2^-R^2^ relationship between AG and SuSiE FM regardless of F-test status (**Fig 4B-C**). Together, these suggest our AG filter performs as well or better than SuSiE finemapping, especially for models which pass the F-test.

**Figure 4:**
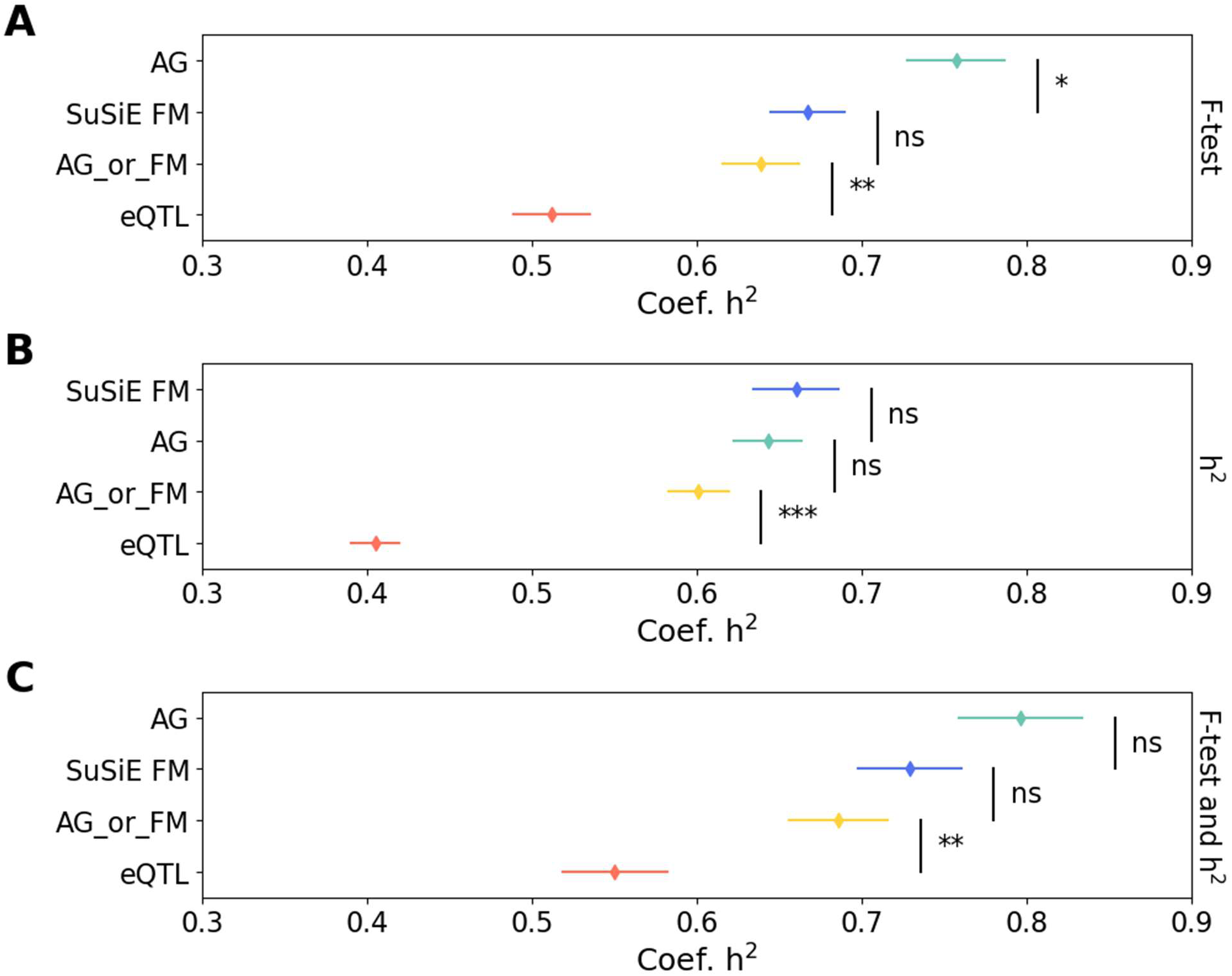
Comparing estimated heritability and EAGLES performance in Geuvadis. EAGLES models were stratified into groups which passed F-test (A), had significant cis-heritability (B) or both (C) based on GTEx whole blood cohort. Forest plots summarize coefficients and 95% confidence intervals for linear models predicting EAGLES model performance in Geuvadis from GCTA cis-heritability (h^2^) estimates in Geuvadis. Permutation test used for pairwise comparisons: ***<0.001; **<0.01; *<0.05; ns>0.05. AG=AlphaGenome; FM=finemapped; AG_or_FM=AlphaGenome or SuSiE finemapped.

AG and FM represent different approaches to nominate potential causal variants, and our previous analyses identified them as the top performing EAGLES eQTL sources. We next directly compared EAGLES performance between these two eQTL sources. Because the genes found in each of our four eQTL sources do not perfectly match, and because F-test outcomes are not always aligned between AG and FM models, we analyzed each gene group separately. Within the F-test only, h^2^ only, and F-test and h^2^ gene groups, we analyzed the subset of the gene group shared by AG and FM, and the subset exclusive to each eQTL source. Across all tissue types, a relatively small fraction of each gene group was represented in both FM and AG eQTL tables, and an even smaller fraction also met F-test or h^2^ criteria for both eQTL sources. More genes met F-test or h^2^ criteria for AG than for FM (**Fig S15-20**). When considering genes exclusive to one eQTL source, EAGLES performance was comparable between AG and FM. However, in comparing models represented by both eQTL sources, FM consistently outperformed AG for a larger number of genes.

### eQTL source and EAGLES performance in hallmark gene sets

Genome-wide approaches like TWAS require large cohorts for gene-trait association testing^34,35^. For smaller cohorts, a hypothesis-driven approach can prioritize a subset of association tests to perform. We used hallmark gene sets from the molecular signatures database (MSigDB^36^) to represent groups of functionally related genes. As the top performing eQTL sources, we compared EAGLES performance between AG and FM models in hallmark gene sets. As EAGLES models were only trained for genes represented in our eQTL tables, we selected the top gene sets for each tissue based on the fraction of gene set members represented among EAGLES models. Because gene representation varied across eQTL sources, we ranked gene sets based on the lowest fraction of members included in each eQTL source, prioritizing gene sets well represented across the four examined eQTL sources (**Fig S21**). Through this, we found example gene sets with relatively high representation across all examined tissues (HALLMARK_DNA_REPAIR), relatively high representation in specific tissue types (HALLMARK_INTERFERON_ALPHA_RESPONSE in whole blood) and with relatively low representation across all examined tissues (HALLMARK_IL6_JAK_STAT3_SIGNALING).

For whole blood, the top 4 gene sets by representation in EAGLES included DNA repair (DNA-R), Interferon-α (IFNα), Interferon-γ (IFNγ) and Reactive Oxygen Species (ROS). Through empirical Cumulative Distribution Function (eCDF) analysis, we found that EAGLES performance varied across these 4 gene sets for FM (**Fig 5A**) and AG (**Fig 5B**). However, for all four of these gene sets, we found that FM models outperformed AG (**Fig 5C-F**) in the subset of genes modeled by both eQTL sources. We observed similar trends across the other five examined tissues with FM models outperforming AG for larger proportions of each hallmark gene set than AG outperformed FM (**Fig S22-26**).

**Figure 5:**
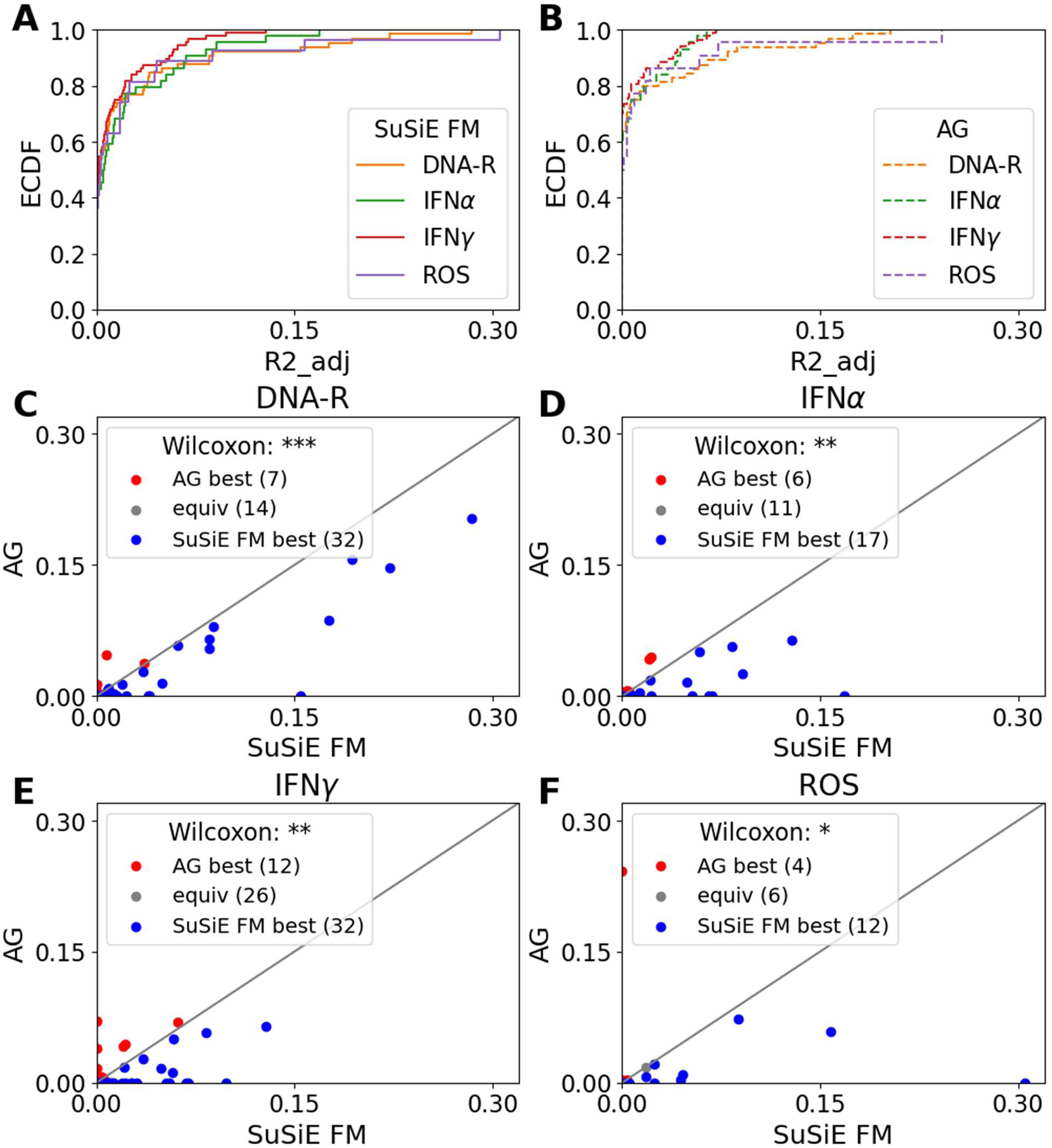
EAGLES performance in hallmark gene sets. ECDF curves show EAGLES FM (A) and AG (B) performance (adjusted R^2^) over 4 hallmark gene sets in the independent blood cohort. For genes represented in both EAGLES FM and EAGLES AG input eQTL tables, scatter plots compare EAGLES FM and AG performance in DNA repair (C), Interferon ɑ (D), Interferon γ (E), and reactive oxygen species (ROS) genes (F). The Wilcoxon signed-rank test was used for comparisons: ***<0.001; **<0.01; *<0.05; ns>0.05. FM=finemapped; AG=AlphaGenome.

### Applications to trait prediction

After identifying gene sets with robust performance, we next asked whether EAGLES scores from these pathways capture germline variation relevant to phenotypic outcomes. The top represented pathways, including IFNα and IFNγ response^37^, ROS signaling^38^, and DNA repair^39^, are all central to immune function and represent good candidates for studying immune phenotypes. As genetic modifiers of immunity have previously been reported to predict immunotherapy response^40,41^, we evaluated whether EAGLES scores for these pathways were informative in predicting response to immune checkpoint blockade (ICB) in melanoma^42^. Further, we sought to determine the extent to which EAGLES modeling choices, such as score construction method and model type, would affect our ability to detect important genes affecting ICB therapy response.

Based on area under the receiver operating characteristic curve (AUC), we saw that EAGLES FM consistently performed best (**Fig 6**) compared to other EAGLES eQTL sources (**Fig S27-29**). As the goal of this analysis was to test whether EAGLES scores carry any detectable germline signal in this prediction task rather than to achieve high predictive accuracy, even modest AUC values were informative. In the logistic classifier, the ROS (AUC=0.68) and DNAR (AUC=0.68) pathways provided the strongest predictive signal for ICB response (**Fig 6A**). In the nonlinear XGBoost classifier, the ROS pathway also yielded a strong signal (AUC=0.70; **Fig 6B**). As the AUC values from the classifiers reflected modest predictive power, we applied Mann-Whitney U tests to assess statistical differences in predicted probabilities between responders and nonresponders. For the ROS pathway, we found better than random performance for the nonlinear classifier over the linear classifier (MWU p-value: logistic=0.050, XGBoost=0.032), highlighting the functionality of nonlinear modeling in this prediction task.

**Figure 6:**
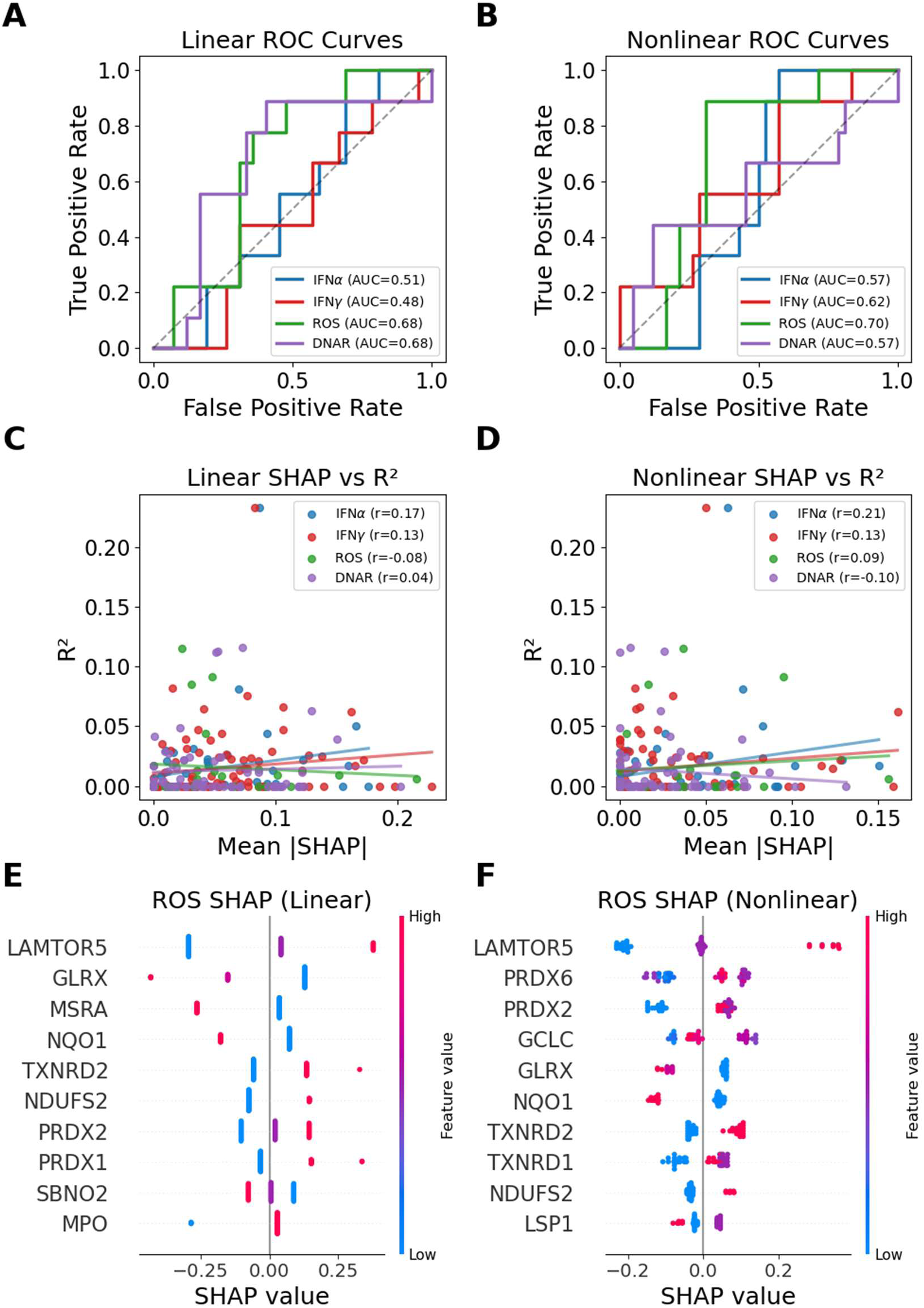
Classifying ICB response from EAGLES finemapped imputed expression. ROC curves and AUC values for logistic classifiers trained on IFNɑ, IFNγ, ROS, and DNAR gene-score features (A) and XGBoost classifiers trained on the same feature sets (B). Scatter plots show relationships between mean absolute SHAP value and EAGLES adjusted R^2^ for gene features in logistic (C) and XGBoost (D) classifiers. SHAP beeswarm plots summarize feature importances in the logistic ROS classifier (E) and in the XGBoost ROS classifier (F). Positive SHAP values push models towards predicting response.

To assess whether underlying genetic architecture reflects predictive utility for a phenotype, we compared each classifier’s feature importance (SHapley Additive exPlanation [SHAP] value^43^) with the feature’s EAGLES performance (adjusted R^2^). This assessment evaluates whether features with higher R^2^ values are also the same genes whose predicted expression contributes more strongly to ICB response. We found weak correlations in both linear (R=-0.08-0.17, **Fig 6C**) and nonlinear (R=-0.10-0.21, **Fig 6D**) classifiers, suggesting that genes with low R^2^ values can still have substantial utility for predicting ICB response.

Finally, we used SHAP values to compare genes contributing to ICB response prediction by the linear (**Fig 6E**) or nonlinear (**Fig 6F**) classifier, focusing on the ROS pathway which showed relatively high performance in both classifier types. Both classifiers consistently highlighted a common set of ROS genes, including *LAMTOR5*, *PRDX2*, *PRDX6*, and *GLRX*, indicating a core biological predictive signal that is stable across classifier types. Notably, the SHAP summary plots showed that the logistic regression models enforce a linear effect of predicted gene expression on response, while XGBoost models appear to learn threshold-based, categorical patterns. For instance, the linear model distributes SHAP values for *PRDX2* equidistantly across different expression levels while the nonlinear model distributes the SHAP values into two rough groups consisting of only low expression predicting nonresponse while intermediate or high expression predicting response. We observed a similar divergence in SHAP distributions for other pathways (**Fig S30**), with nonlinear models showing more finer structured SHAP patterns compared to the linear classifiers. Further, we saw that the different classifiers also prioritized different gene-level features for making predictions, providing a potential explanation for the improved predicted performance of nonlinear classifiers.

## Discussion

EAGLES is a versatile method for constructing QTL-based gene scores, implemented in Nextflow and designed for ease of use. Though we focused on eQTLs in the present study, EAGLES pipelines are flexible to QTL type, so other QTL sources such as the Human Kidney meQTL Atlas^44^ or the eQTM Atlas^45^ can also be used to train EAGLES models. Unlike existing tools like FUSION and PrediXcan, which are limited to linear poly-eQTL sources, EAGLES supports both linear and nonlinear model types, enabling better performance when gene expression is not well captured by linear effects. In fact, we found multiple tissue contexts where the nonlinear XGBoost model type performed best (Figure S7). While elastic net generally outperformed other model types, we found multiple tissue contexts where it was outperformed by nonlinear models.

When applying EAGLES-derived scores to a trait prediction task, we found that classifier type can impact the detection of trait-relevant genes. By using gene scores constructed from finemapped eQTLs as features with both linear and nonlinear classifiers, we showed that EAGLES scores can recapitulate prior work linking the ROS pathway to ICB therapy response^38^, demonstrating that these scores capture modest but detectable germline signal. Notably, we detected stronger associations when using a nonlinear XGBoost classifier, suggesting nonlinear classifiers may be advantageous for traits shaped by complex interactions between features.

The quality of gene scores produced by tools like EAGLES depends on the quality of input eQTLs. We found finemapping methods like SuSiE and sequence to function models like AlphaGenome can help refine eQTL sets, improving expression prediction performance. It is known that the impacts of eQTLs on gene regulation and disease states can be cell type specific^4,5,46^, suggesting that cell-based models may be useful for traits with known causal cell types. In practice, EAGLES can be used to train single-cell (sc) based models if sc-eQTLs have been identified. Recent efforts like sc-eQTLGen^47^, CIMA^48^ and OneK1K^49^ have focused on immune cell populations, with more cell and tissue types anticipated in future sc-eQTL studies. In the present study, we focused our evaluation of EAGLES models on multi-tissue cohorts with bulk RNA-seq, but future benchmarks of EAGLES parameter choices in single-cell QTL-derived models will be useful.

Cross ancestry portability of eQTL-based scores remains a challenging problem^50,51^. To evaluate EAGLES performance in independent cohorts, we restricted our analysis to European ancestry to maximize sample sizes in large multi-tissue cohorts GTEx and TCGA. Though not evaluated here, we expect EAGLES models will perform best within the same ancestry group used for eQTL discovery. Cohorts like MAGE^52^, SAGE II^53^ and GALA II^54^ have been used for eQTL discovery in non-European individuals^52,55^, and would be valuable in training genetically diverse EAGLES models. As these and other^56–58^ efforts to identify eQTLs in non-European ancestry groups have largely focused on whole blood or lymphoblastoid cell lines, the utility of EAGLES outside European ancestry groups can be improved by future eQTL studies with both tissue and genetic diversity.

## Conclusions

We developed the EAGLES pipelines to facilitate eQTL-based gene expression model training and evaluation. We evaluated parameter choices by training models using GTEx, and benchmarking performance in independent cohorts (TCGA or Geuvadis). Using ICB therapy response as an example trait, we demonstrated that EAGLES scores can be used to identify trait-relevant genes and pathways.

## Methods

### Genotype Curation

GTEx genotypes were downloaded from dbGaP (phs000424.v8). We performed quality control on the genotypes following the steps from Syed et al. 2025^59^ (see Plink-QC below) and filtered the result to minor allele frequency (MAF) >= 1% for subsequent analyses.

TCGA genotypes from Affymetrix SNP 6.0 were acquired from Genomic Data Commons^60^ and were then lifted over to GRCh38, phased, and imputed using the TOPMed imputation server^61^. We performed quality control on the imputed genotypes (see Plink-QC below) and filtered the result to minor allele frequency (MAF) >= 1% for subsequent analyses.

Geuvadis samples were derived as a subset of the 1000 Genomes Project (1kGP) Consortium^62^. Phased, chromosome level call files were concatenated and normalized with standard procedures including the decomposition of multiallelic variants into biallelic variants, variant atomization, and left-alignment to the GRCh38 reference genome. Geuvadis samples were selected using sample identifiers from recount3 metadata (accession ERP001942), and the subset was extracted from the full 1kGP dataset using PLINK2^30^. ICB cohort genotypes were processed as previously described^40,41^.

### Genotype pre-processing

We performed quality control on genotype data from GTEx using a custom pipeline implemented in Nextflow based on plinkQC^59^. For each chromosome, various QC metrics were calculated for variants including missingness, allele counts, and Hardy Weinberg equilibrium (HWE). Samples were discarded if deviating more than 3 standard deviations from the mean or with >3% missing genotypes. Variants were discarded with a stringent missingness threshold (F_MISS < 1 × 10^-7^) and HWE deviation (p > 0.01). Next, variants were linkage disequilibrium (LD)-pruned using a sliding window approach (50 variants per window, step size of 5 variants, and pairwise r^2^ threshold = 0.7) after excluding high-LD regions. These variants were then used to estimate pairwise relatedness via identity by descent (IBD) using the KING method (flagging pairs with PI_HAT > 0.177, suggesting first-degree relatives). No samples exceeded this threshold, and therefore no samples were excluded based on relatedness. We also assessed population structure with principal component analysis (PCA) after merging the GTEx samples with the 1kGP reference panel. Variants and samples passing all QC filters across chromosomes were aggregated, and the final QC-ed dataset was created by subsetting to the union of passing variants and samples.

In order to ensure consistent variant representation across our cohorts, we utilized GRIEVOUS^63^, a command-line tool designed for multi-dataset variant harmonization. All datasets were first converted to PLINK2 format and restricted to biallelic SNPs. To ensure compatibility with GRIEVOUS indexing requirements, variant identifiers were standardized to a consistent format (CHROM:POS:REF:ALT) using PLINK2. The datasets were further partitioned by chromosome. We selected GTEx for initialization of the GRIEVOUS database which defines the reference orientation of variants. Chromosome level variant files were processed with the *realign* workflow which removes any duplicate or invalid variants, registers the variants into a unified index, and sets the allele orientation relative to the internal database. The output of this command results in a report specifying which variants required reorientation. We subsequently processed TCGA and Geuvadis to the pre-initialized GTEx-based GRIEVOUS database. For each dataset, we used PLINK2 to set reference and alternative allele definitions. Finally, chromosome level outputs were merged back together to complete aligned datasets.

### Training, held-out and independent cohort filtering

GRAF-pop^64^ was used to assign local ancestry to all samples across cohorts, and EUR samples were retained. We split EUR individuals in the GTEx cohort, using 70% for training and the remaining 30% of individuals as a held out test set. EUR individuals from Geuvadis and TCGA were used as independent cohorts.

### Statistics

We used Ezekiel’s adjusted R^2^ statistic^65^ as the adjusted R^2^ metric used to quantify EAGLES gene expression prediction performance. We used the Scipy^66^ implementations of the Wilcoxon signed-rank test (single gene group) and the Mann-Whitney U test (multiple gene groups) to compare prediction performance between different EAGLES modes.

We applied a permutation test to compare differences in adjusted R^2^ vs. h^2^ slope between EAGLES eQTL sources. For 1000 permutations, we first pooled R^2^ and h^2^ values from a pair of eQTL sources, shuffled them, and randomly partitioned them into two groups. Then we fit separate linear regression of R^2^ on h^2^ for the two permuted groups, and recorded the difference in slopes (**β**_i,2_-**β**_i,1_; for *i*th permutation) to construct a background null distribution. For the eQTL sources A and B, we computed the difference in slope (**β**_B_ - **β**_A_). The p-value for testing **β**_B_ > **β**_A_ was computed as the fraction of **β**_i,2_ - **β**_i,1_ values greater than **β**_B_ - **β**_A._

### The EAGLES Pipelines

#### EAGLES eQTL Tables

For each tissue, the GTEx v8 EUR eQTL tables were downloaded from the GTEx portal and used for EAGLES eQTL models. The GTEx v10 SuSiE finemapped eQTL tables were downloaded from the GTEx portal, and variants without annotated aFC values were removed to leave a single eQTL from each credible set. SuSiE finemapped tables were used for EAGLES FM tables.

The GTEx eQTL tables were filtered to prioritize putatively functional variants using AlphaGenome^29^. For each variant, functional impact scores were computed with AlphaGenome’s score_variant function across three modalities - RNA-seq expression, chromatin accessibility (DNase-seq), and transcription start site initiation (CAGE). For the RNA-seq track scoring was GTEx tissue-matched, and for the DNase and CAGE tracks, scoring was restricted to biosamples corresponding to relevant cell and tissue types (ex: blood-derived cell populations for whole blood variant scoring). AlphaGenome scores were summarized per variant by taking the maximum absolute effect across the tracks. Because raw scores from different modalities are on different scales, we used AlphaGenome’s quantile_score associated with each variant, which ranks the variant’s score relative to a background distribution of common variants. We deemed variants to be functionally important if their absolute quantile score was > 0.9 in at least one of the three modalities. These AlphaGenome filtered tables were used for EAGLES AG models.

We constructed AG_or_FM eQTLs tables as the union of AG and FM tables for each tissue, and used these for EAGLES AG_or_FM models.

#### Expression outlier filtering

As previously defined^67^, for each gene we excluded outlier samples with expression > 5TPM and 5 inter-quartile range (IQR) above the 75th percentile.

#### LD filtering

We used PLINK2^30^ to extract cis-eQTLs defined by the input eQTL table and to apply specified LD pruning (*LDNone*: no pruning, *LDLax*: 100kb window and 0.8 threshold, *LDMed*: 200kb window and 0.5 threshold, *LDStrict*: 500kb window and 0.2 threshold).

#### Model Types

We implemented EAGLES Flip Allele Count (FA) as a linear model. From input eQTLs, model coefficients are set to +1 or −1, consistent with the sign of the eQTL slope. To ensure only positive scores are returned, the model intercept is set to 2* C_neg_ where C_neg_ is the number of all negative coefficients.

We implemented EAGLES principal component regression (PCR) as follows. The PC matrix X* was computed as

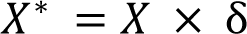

where X are genotypes and **δ** are the principal component loadings. Consistent with MAGMA^22^ default settings, PCs accounting for bottom 0.1% variation are pruned. Then expression (Y) of a gene is modeled as

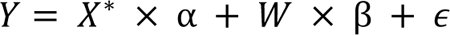

where X* are PCs computed over cis-eQTLs associated with the modeled gene, W are covariates, vector α represents the genetic effect, vector β the effect of the covariates, and ɛ represents residuals. After fitting this PC-based model using SciKit-Learn^68^ LinearRegression, the final EAGLES SNP-based model is represented as

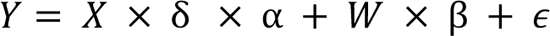

We implemented EAGLES elastic net (EN) using SciKit-Learn ElasticNetCV with L1 ratio of 0.5, 5-folder cross-validation and maximum iteration of 10000.

We implemented EAGLES XGBoost (XGB) using the XGBoost package^69^. We used Optuna^70^ for hyperparameter tuning, running 200 trials and a 5-fold cross validation procedure.

In the special case that a model was trained from a single feature, a SciKit-Learn linear regression model was trained regardless of EAGLES mode.

#### Scoring EAGLES Models

Given a list of trained EAGLES models, PLINK2 files from the cohort to be scored and covariates from the cohort to be scored, the scoring pipeline produced four output files summarizing model performances. First, the EAGLES predictions from each model were returned in the scores output file. Second, a summary of the number of total features per model and the number of missing features from the PLINK2 and covariate files of the scored cohort were returned in the missing_counts output file. Third, the mean absolute SHAP^43^ values for each feature in each scored model were returned in the feature_summary output file. Features with one unique value in the scored cohort or a mean absolute SHAP value of 0 were excluded from this output. Finally, a summary of each model’s performance was returned in the model_performance output file.

### ICB therapy response classification

For classifier training, we used three independent melanoma ICB cohorts (Hugo et al.^71^, Snyder et al.^72^ and Riaz et al.^73^), and evaluated performance in an independent melanoma cohort (Van Allen et al.^74^). ICB response was defined as either complete response, partial response, or stable disease.

We trained both logistic regression models and XGBoost models to predict immunotherapy response. For the logistic regression models, all features were scaled to the [0,1] range with SciKit-Learn min-max normalization by first fitting the scaler on the training data only and then applying the scaler to the test data. Logistic regression models were L2-regularized and optimized with LBFGS with a maximum of 5,000 iterations. XGBoost models were trained with a binary logistic objective with regularized, shallow trees and 300 boosting rounds.

To evaluate classifier performance, predicted probabilities for each hallmark gene set were obtained from XGBoost on the held out test data. With these probabilities we computed the area under the ROC curves (auROC), and tested for statistical significance using the one-sided Mann-Whitney U test to investigate whether classifiers assigned higher predicted probabilities to true responders compared to nonresponders. To evaluate the feature importance of the classifiers, we computed SHAP^43^ values to assess the contribution of each gene score on the predicted probability of response.

## Declarations

### Ethics approval and consent to participate

Not applicable

### Consent for publication

Not applicable

### Availability of data and materials

The EAGLES Nextflow pipeline, along with scripts used to preprocess and QC PLINK2 files for this study, can be found at https://github.com/cartercompbio/EAGLES (Zenodo accession: https://doi.org/10.5281/zenodo.21983535).

Pretrained models used in the current study are available through Zenodo at https://doi.org/10.5281/zenodo.21477399.

## Authors’ contributions

D.M built the EAGLES pipelines, carried out analyses, interpreted the results, and wrote the manuscript. N.P assisted with building the EAGLES pipelines, curated and preprocessed data from Geuvadis and GTEx cohorts, implemented the plinkqc pipeline and assisted with manuscript preparation. D.L curated and preprocessed data from TCGA cohorts, assisted with curation and preprocessing of the Geuvadis cohort, assisted with the plinkqc pipeline implementation, and assisted with manuscript preparation. P.S imputed germline data from the TCGA cohort. T.A. assisted with manuscript preparation and revision. L.B.A supervised the project. H.K designed and supervised the project, and wrote the manuscript. All authors read and approved the final manuscript.

## Acknowledgements

We thank Dr. Ammal Abbasi, Jessica Au, Dr. Mariya Kazachkova, and Dr. Christopher Steele for their helpful comments. We thank Dr. Meghana Pagadala for assistance in preprocessing germline variants from the TCGA cohort.

## References

1. Gong, J. et al. PancanQTL: systematic identification of cis-eQTLs and trans-eQTLs in 33 cancer types. Nucleic Acids Res. 46, D971–D976 (2018).

2. The GTEx Consortium. The GTEx Consortium atlas of genetic regulatory effects across human tissues. Science 369, 1318–1330 (2020).

3. Díez-Obrero, V. et al. Genetic Effects on Transcriptome Profiles in Colon Epithelium Provide Functional Insights for Genetic Risk Loci. Cell. Mol. Gastroenterol. Hepatol. 12, 181–197 (2021).

4. Yazar, S. et al. Single-cell eQTL mapping identifies cell type–specific genetic control of autoimmune disease. Science 376, eabf3041 (2022).

5. Fu, Y. et al. Single-cell eQTL mapping reveals cell-type-specific genetic regulation in lung cancer. Cell Genomics 0, (2025).

6. Wang, C. et al. Cell type-specific eQTL analysis of COVID-19 based on single-cell transcriptomic data. NAR Genomics Bioinforma. 7, lqaf130 (2025).

7. Ding, R. et al. scQTLbase: an integrated human single-cell eQTL database. Nucleic Acids Res. 52, D1010–D1017 (2024).

8. Chen, C. et al. Single-cell eQTL Mapping Reveals Cell Subtype–specific Genetic Control and Mechanism in Malignant Transformation of Colorectal Cancer. Cancer Discov. 15, 1649–1675 (2025).

9. Hao, K. et al. Lung eQTLs to Help Reveal the Molecular Underpinnings of Asthma. PLOS Genet. 8, e1003029 (2012).

10. Li, Q. et al. Expression QTL-based analyses reveal candidate causal genes and loci across five tumor types. Hum. Mol. Genet. 23, 5294–5302 (2014).

11. Miao, X.-C. et al. Cell-type-specific cis-eQTLs in pancreatic cell types identify novel risk genes for type 2 diabetes. Brief. Bioinform. 26, bbaf531 (2025).

12. Nica, A. C. & Dermitzakis, E. T. Expression quantitative trait loci: present and future. Philos. Trans. R. Soc. B Biol. Sci. 368, 20120362 (2013).

13. Sudlow, C. et al. UK Biobank: An Open Access Resource for Identifying the Causes of a Wide Range of Complex Diseases of Middle and Old Age. PLOS Med. 12, e1001779 (2015).

14. Bick, A. G. et al. Genomic data in the All of Us Research Program. Nature 627, 340–346 (2024).

15. Le Guen, Y. et al. eQTL of KCNK2 regionally influences the brain sulcal widening: evidence from 15,597 UK Biobank participants with neuroimaging data. Brain Struct. Funct. 224, 847–857 (2019).

16. Kerimov, N. et al. eQTL Catalogue 2023: New datasets, X chromosome QTLs, and improved detection and visualisation of transcript-level QTLs. PLOS Genet. 19, e1010932 (2023).

17. Võsa, U. et al. Large-scale cis- and trans-eQTL analyses identify thousands of genetic loci and polygenic scores that regulate blood gene expression. Nat. Genet. 53, 1300–1310 (2021).

18. Lonsdale, J. et al. The Genotype-Tissue Expression (GTEx) project. Nat. Genet. 45, 580– 585 (2013).

19. The GTEx Consortium. The GTEx Consortium atlas of genetic regulatory effects across human tissues. Science 369, 1318–1330 (2020).

20. Gamazon, E. R. et al. A gene-based association method for mapping traits using reference transcriptome data. Nat. Genet. 47, 1091–1098 (2015).

21. Gusev, A. et al. Integrative approaches for large-scale transcriptome-wide association studies. Nat. Genet. 48, 245–252 (2016).

22. de Leeuw, C. A., Mooij, J. M., Heskes, T. & Posthuma, D. MAGMA: Generalized Gene-Set Analysis of GWAS Data. PLOS Comput. Biol. 11, e1004219 (2015).

23. Gerring, Z. F., Mina-Vargas, A., Gamazon, E. R. & Derks, E. M. E-MAGMA: an eQTL-informed method to identify risk genes using genome-wide association study summary statistics. Bioinformatics 37, 2245–2249 (2021).

24. Grinberg, N. F. & Wallace, C. Multi-tissue transcriptome-wide association studies. Genet. Epidemiol. 45, 324–337 (2021).

25. Okoro, P. C. et al. Transcriptome prediction performance across machine learning models and diverse ancestries. Hum. Genet. Genomics Adv. 2, 100019 (2021).

26. Wang, G., Sarkar, A., Carbonetto, P. & Stephens, M. A Simple New Approach to Variable Selection in Regression, with Application to Genetic Fine Mapping. J. R. Stat. Soc. Ser. B Stat. Methodol. 82, 1273–1300 (2020).

27. Wen, X., Lee, Y., Luca, F. & Pique-Regi, R. Efficient Integrative Multi-SNP Association Analysis via Deterministic Approximation of Posteriors. Am. J. Hum. Genet. 98, 1114–1129 (2016).

28. Barbeira, A. N. et al. Fine-mapping and QTL tissue-sharing information improves the reliability of causal gene identification. Genet. Epidemiol. 44, 854–867 (2020).

29. Avsec, Ž., et al. Advancing regulatory variant effect prediction with AlphaGenome. Nature 649, 1206–1218 (2026).

30. Chang, C. C. et al. Second-generation PLINK: rising to the challenge of larger and richer datasets. GigaScience 4, s13742–015-0047–8 (2015).

31. Lappalainen, T. et al. Transcriptome and genome sequencing uncovers functional variation in humans. Nature 501, 506–511 (2013).

32. Weinstein, J. N. et al. The Cancer Genome Atlas Pan-Cancer analysis project. Nat. Genet. 45, 1113–1120 (2013).

33. Liu, S., Lu, M., Li, H. & Zuo, Y. Prediction of Gene Expression Patterns With Generalized Linear Regression Model. Front. Genet. 10, (2019).

34. He, R., Xue, H., Pan, W. & Initiative, for the A. D. N. Statistical power of transcriptome-wide association studies. Genet. Epidemiol. 46, 572–588 (2022).

35. Baranger, D. A. A. et al. Multi-omics cannot replace sample size in genome-wide association studies. Genes Brain Behav. 22, e12846 (2023).

36. Liberzon, A. et al. The Molecular Signatures Database (MSigDB) hallmark gene set collection. Cell Syst. 1, 417–425 (2015).

37. Belardelli, F. Role of interferons and other cytokines in the regulation of the immune response. APMIS 103, 161–179 (1995).

38. Yang, Y., Bazhin, A. V., Werner, J. & Karakhanova, S. Reactive Oxygen Species in the Immune System. Int. Rev. Immunol. 32, 249–270 (2013).

39. Nakad, R. & Schumacher, B. DNA Damage Response and Immune Defense: Links and Mechanisms. Front. Genet. 7, (2016).

40. Pagadala, M. et al. Germline modifiers of the tumor immune microenvironment implicate drivers of cancer risk and immunotherapy response. Nat. Commun. 14, 2744 (2023).

41. Sears, T. J. et al. Integrated Germline and Somatic Features Reveal Divergent Immune Pathways Driving Response to Immune Checkpoint Blockade. Cancer Immunol. Res. 12, 1780–1795 (2024).

42. Haanen, J. B. A. G. Immunotherapy of melanoma. Eur. J. Cancer Suppl. 11, 97–105 (2013).

43. Lundberg, S. M. & Lee, S.-I. A Unified Approach to Interpreting Model Predictions. in Advances in Neural Information Processing Systems vol. 30 (Curran Associates, Inc., 2017).

44. Liu, H. et al. Epigenomic and transcriptomic analyses define core cell types, genes and targetable mechanisms for kidney disease. Nat. Genet. 54, 950–962 (2022).

45. Sriram, A. et al. eQTM (expression quantitative trait methylation) Atlas: a comprehensive resource of over 11 million DNA methylation–gene expression associations through across 11 tissues and 4 diseases. 2026.06.07.730721 Preprint at 10.64898/2026.06.07.730721 (2026).

46. Natri, H. M. et al. Cell-type-specific and disease-associated expression quantitative trait loci in the human lung. Nat. Genet. 56, 595–604 (2024).

47. Kaptijn, D. et al. Federated single-cell QTL meta-analysis reveals novel disease mechanisms. 2026.01.20.700519 Preprint at 10.64898/2026.01.20.700519 (2026).

48. Yin, J. et al. Chinese Immune Multi-Omics Atlas. Science 391, eadt3130 (2026).

49. Yazar, S. et al. Single-cell eQTL mapping identifies cell type–specific genetic control of autoimmune disease. Science 376, eabf3041 (2022).

50. Mogil, L. S. et al. Genetic architecture of gene expression traits across diverse populations. PLOS Genet. 14, e1007586 (2018).

51. Kachuri, L. et al. Gene expression in African Americans, Puerto Ricans and Mexican Americans reveals ancestry-specific patterns of genetic architecture. Nat. Genet. 55, 952– 963 (2023).

52. Taylor, D. J. et al. Sources of gene expression variation in a globally diverse human cohort. Nature 632, 122–130 (2024).

53. White, M. J. et al. Novel genetic risk factors for asthma in African American children: Precision Medicine and the SAGE II Study. Immunogenetics 68, 391–400 (2016).

54. Oh, S. S. et al. Effect of secondhand smoke on asthma control among black and Latino children. J. Allergy Clin. Immunol. 129, 1478–1483.e7 (2012).

55. Kachuri, L. et al. Gene expression in African Americans, Puerto Ricans and Mexican Americans reveals ancestry-specific patterns of genetic architecture. Nat. Genet. 55, 952– 963 (2023).

56. Chen, W. et al. Expression Quantitative Trait Loci (eQTL) Mapping in Puerto Rican Children. PLOS ONE 10, e0122464 (2015).

57. Shang, L. et al. Genetic Architecture of Gene Expression in European and African Americans: An eQTL Mapping Study in GENOA. Am. J. Hum. Genet. 106, 496–512 (2020).

58. Wen, J. et al. Gene expression and splicing QTL analysis of blood cells in African American participants from the Jackson Heart Study. Genetics 228, iyae098 (2024).

59. Syed, M., Walter, C. & Meyer, H. V. plinkQC: An Integrated Tool for Ancestry Inference, Sample Selection, and Quality Control in Population Genetics. BioRxiv Prepr. Serv. Biol. 2025.11.25.690541 (2025) doi:10.1101/2025.11.25.690541.

60. Heath, A. P. et al. The NCI Genomic Data Commons. Nat. Genet. 53, 257–262 (2021).

61. Das, S. et al. Next-generation genotype imputation service and methods. Nat. Genet. 48, 1284–1287 (2016).

62. Auton, A. et al. A global reference for human genetic variation. Nature 526, 68–74 (2015).

63. Talwar, J. V., Klie, A., Pagadala, M. S. & Carter, H. GRIEVOUS: your command-line general for resolving cross-dataset genotype inconsistencies. Bioinformatics 40, btae489 (2024).

64. Jin, Y., Schaffer, A. A., Feolo, M., Holmes, J. B. & Kattman, B. L. GRAF-pop: A Fast Distance-Based Method To Infer Subject Ancestry from Multiple Genotype Datasets Without Principal Components Analysis. G3 GenesGenomesGenetics 9, 2447–2461 (2019).

65. Hittner, J. B. Ezekiel’s classic estimator of the population squared multiple correlation coefficient: Monte Carlo-based extensions and refinements. J. Gen. Psychol. 147, 213– 227 (2020).

66. Virtanen, P. et al. SciPy 1.0: fundamental algorithms for scientific computing in Python. Nat. Methods 17, 261–272 (2020).

67. Xie, C. et al. Patterns of extreme outlier gene expression suggest an edge of chaos effect in transcriptomic networks. Genome Biol. 26, 272 (2025).

68. Pedregosa, F. et al. Scikit-learn: Machine Learning in Python. Mach. Learn. PYTHON.

69. Chen, T. & Guestrin, C. XGBoost: A Scalable Tree Boosting System. in *Proceedings of the 22nd ACM SIGKDD International Conference on Knowledge Discovery and Data Mining* 785–794 (Association for Computing Machinery, New York, NY, USA, 2016). doi:10.1145/2939672.2939785.

70. Akiba, T., Sano, S., Yanase, T., Ohta, T. & Koyama, M. Optuna: A Next-generation Hyperparameter Optimization Framework. Preprint at 10.48550/arXiv.1907.10902 (2019).

71. Hugo, W. et al. Genomic and Transcriptomic Features of Response to Anti-PD-1 Therapy in Metastatic Melanoma. Cell 168, 542 (2017).

72. Snyder, A. et al. Genetic Basis for Clinical Response to CTLA-4 Blockade in Melanoma. N. Engl. J. Med. 371, 2189–2199 (2014).

73. Riaz, N. et al. Tumor and Microenvironment Evolution during Immunotherapy with Nivolumab. Cell 171, 934–949.e16 (2017).

74. Van Allen, E. M. et al. Genomic correlates of response to CTLA-4 blockade in metastatic melanoma. Science 350, 207–211 (2015).

