## Supplemental Figures and Tables for "Evaluating Aggregated Gene Level eQTL Scores"

Table S1: Number of samples per tissue in training, held-out and independent cohorts

| Tissue | Independent Cohort Name | GTEx Training | GTEx Held-out | Independent |
|---|---|---|---|---|
| breast | TCGA | 231 | 96 | 103 |
| colonSigmoid | TCGA | 186 | 77 | 47 |
| colonTransverse | TCGA | 208 | 84 | 47 |
| lung | TCGA | 307 | 127 | 99 |
| thyroid | TCGA | 333 | 145 | 44 |
| wholeBlood | Geuvadis | 387 | 169 | 362 |

Values indicate the number of samples in the training and held-out partitions of the GTEx cohort, and in the independent cohorts (Geuvadis for blood; TCGA for other tissues).

Table S2: Total number of EAGLES models trained and evaluated

| eQTL Source | LD Threshold | breast | colonSigmoid | colonTransverse | lung | thyroid | wholeBlood |
|---|---|---|---|---|---|---|---|
| eqtl | LDLax | 40765 | 38870 | 43301 | 53218 | 67118 | 45760 |
| eqtl | LDMed | 40902 | 39012 | 43501 | 53490 | 67399 | 45947 |
| eqtl | LDStrict | 41021 | 39147 | 43757 | 53650 | 67628 | 46159 |
| AG | LDNone | 29555 | 31474 | 34816 | 41956 | 63175 | 32459 |
| AG | LDLax | 29487 | 31417 | 34700 | 41874 | 62591 | 32387 |
| AG | LDMed | 29624 | 31552 | 34884 | 42009 | 62911 | 32473 |
| AG | LDStrict | 29667 | 31639 | 34965 | 42079 | 63150 | 32520 |
| FM | LDNone | 26396 | 25963 | 27459 | 32608 | 42108 | 31208 |
| FM | LDLax | 25584 | 25040 | 26640 | 31532 | 40240 | 30028 |
| AG_or_FM | LDNone | 34039 | 34152 | 37498 | 45690 | 63534 | 38782 |
| AG_or_FM | LDLax | 33817 | 33884 | 37277 | 45314 | 62908 | 38315 |
| AG_or_FM | LDMed | 34003 | 34100 | 37446 | 45596 | 63199 | 38603 |
| AG_or_FM | LDStrict | 34103 | 34257 | 37576 | 45743 | 63486 | 38760 |

For each eQTL table, LD threshold, and tissue type, values indicate the number of EAGLES models that were trained and evaluated.

Figure S1

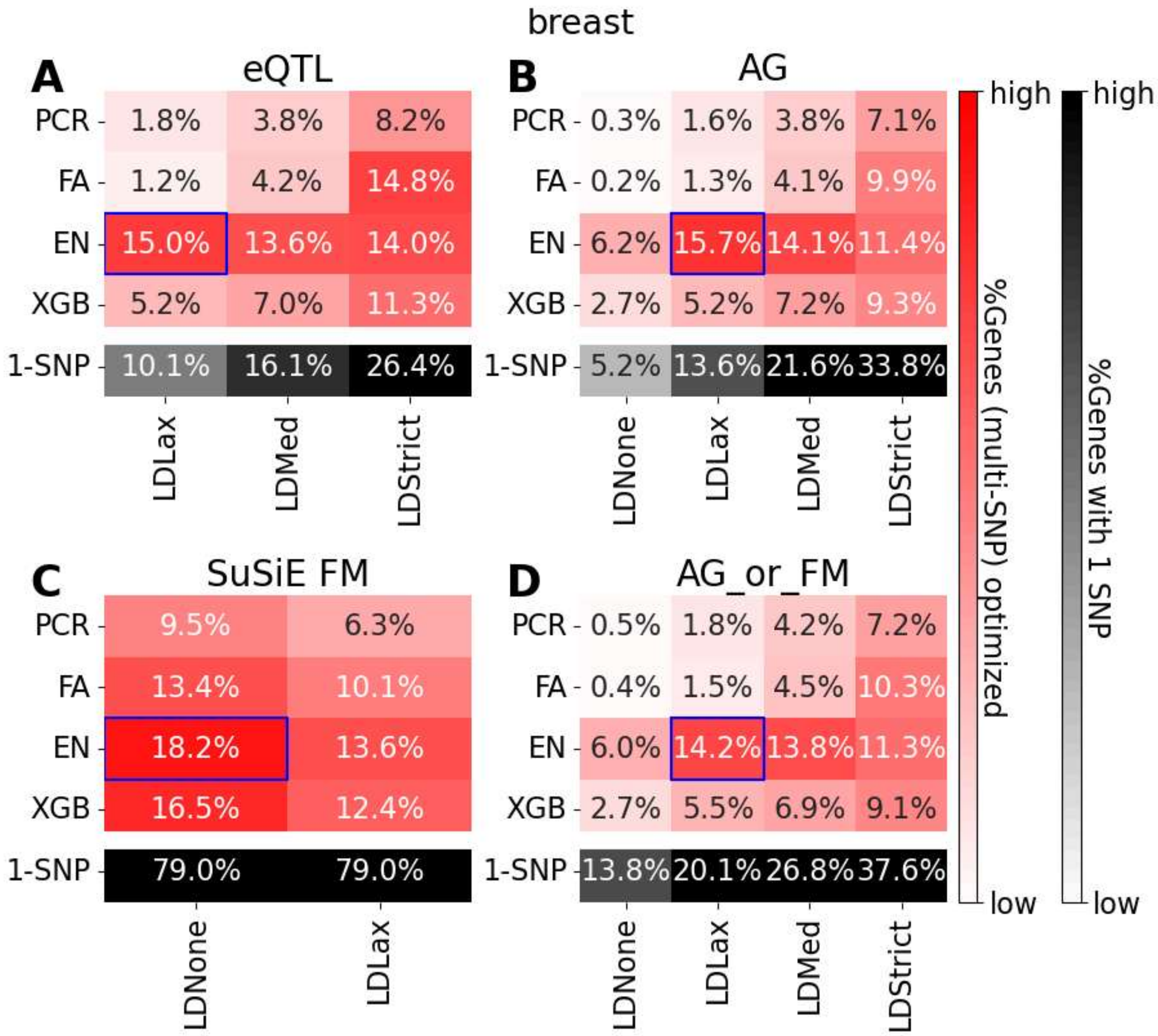


Summary of EAGLES modes impact on adjusted $R^2$ in held-out GTEx breast samples. EAGLES models were trained using GTEx eQTLs (A), AlphaGenome filtered eQTLs (B), GTEx finemapped eQTLs (C) or the union of AlphaGenome and finemapped eQTLs (D). Heatmaps show the fraction of polygenic EAGLES models which maximize performance (adjusted $R^2$ ) relative to other tested EAGLES modes. The black row indicates the fraction of models with a single SNP. Blue boxes indicate the best EAGLES mode. PCR=principal component regression, FA=flip allele count, EN=elasticnet, XGB=xgboost, LD=linkage disequilibrium, FM=finemapped, AG=AlphaGenome.

Figure S2

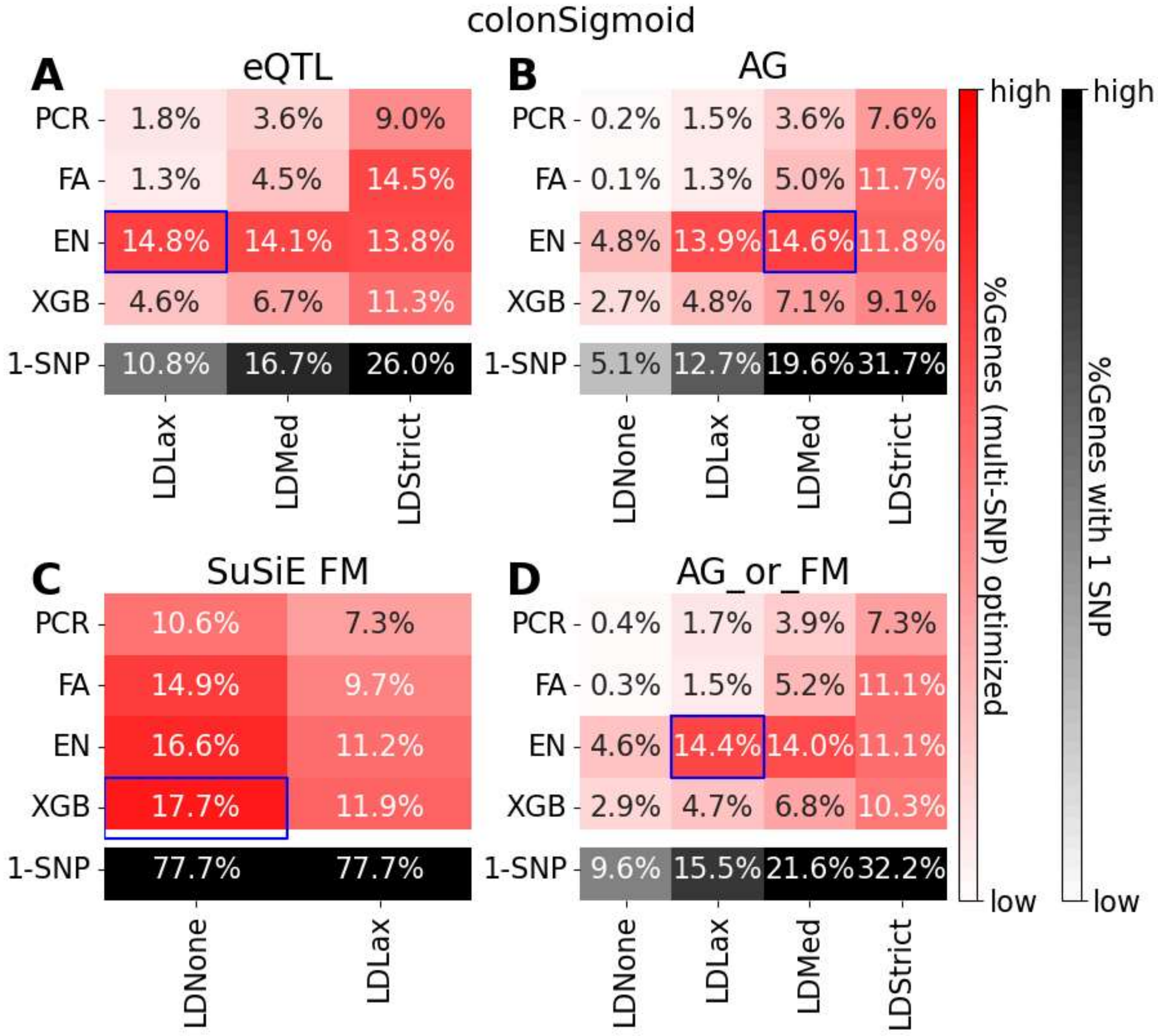


Summary of EAGLES modes impact on adjusted $R^2$ in held-out GTEx sigmoid colon samples. EAGLES models were trained using GTEx eQTLs (A), AlphaGenome filtered eQTLs (B), GTEx finemapped eQTLs (C) or the union of AlphaGenome and finemapped eQTLs (D). Heatmaps show the fraction of polygenic EAGLES models which maximize performance (adjusted $R^2$ ) relative to other tested EAGLES modes. The black row indicates the fraction of models with a single SNP. Blue boxes indicate the best EAGLES mode. PCR=principal component regression, FA=flip allele count, EN=elasticnet, XGB=xgboost, LD=linkage disequilibrium, FM=finemapped, AG=AlphaGenome.

Figure S3

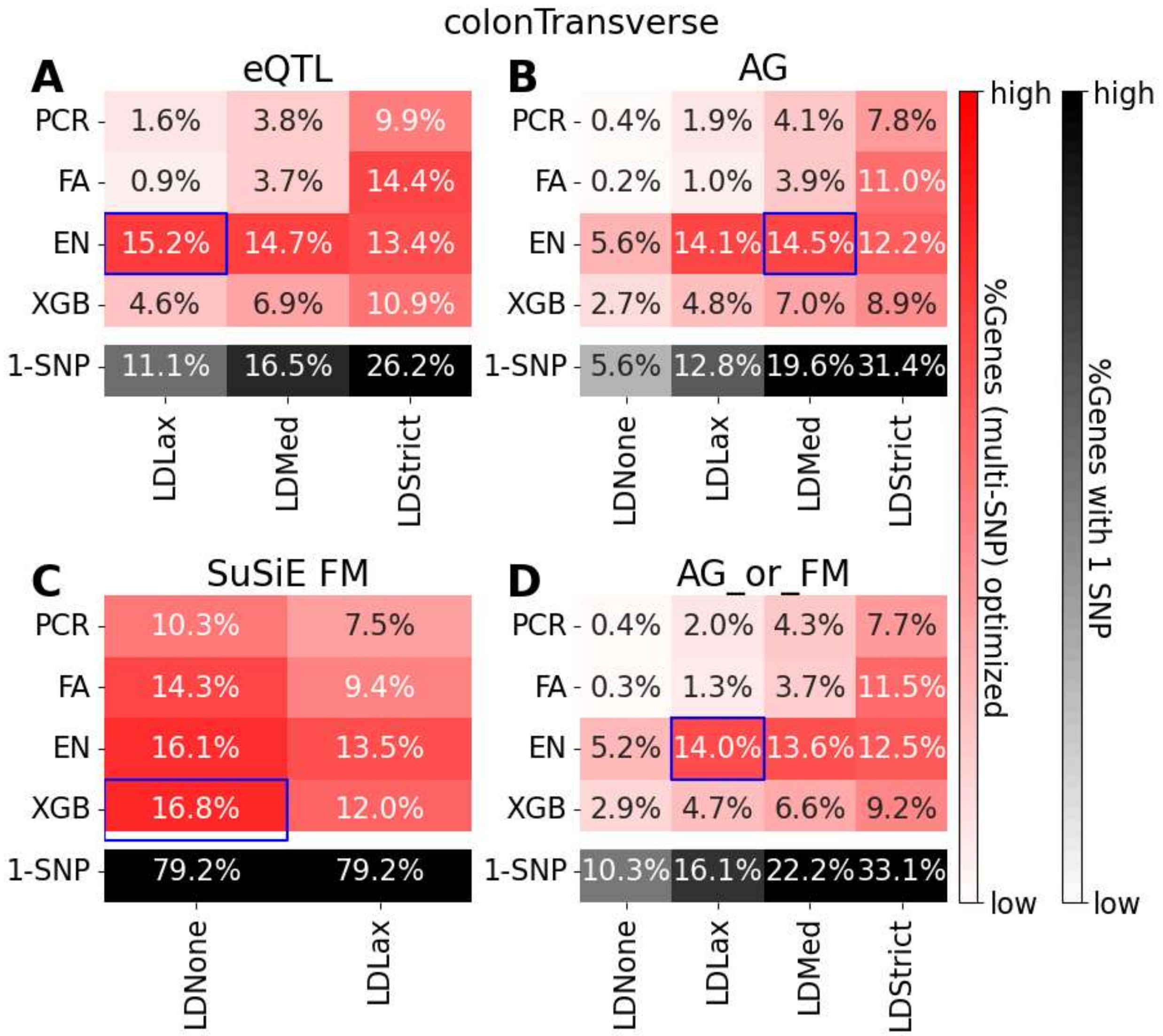


Summary of EAGLES modes impact on adjusted $R^2$ in held-out GTEx transverse colon samples. EAGLES models were trained using GTEx eQTLs (A), AlphaGenome filtered eQTLs (B), GTEx finemapped eQTLs (C) or the union of AlphaGenome and finemapped eQTLs (D). Heatmaps show the fraction of polygenic EAGLES models which maximize performance (adjusted $R^2$ ) relative to other tested EAGLES modes. The black row indicates the fraction of models with a single SNP. Blue boxes indicate the best EAGLES mode. PCR=principal component regression, FA=flip allele count, EN=elasticnet, XGB=xgboost, LD=linkage disequilibrium, FM=finemapped, AG=AlphaGenome.

Figure S4

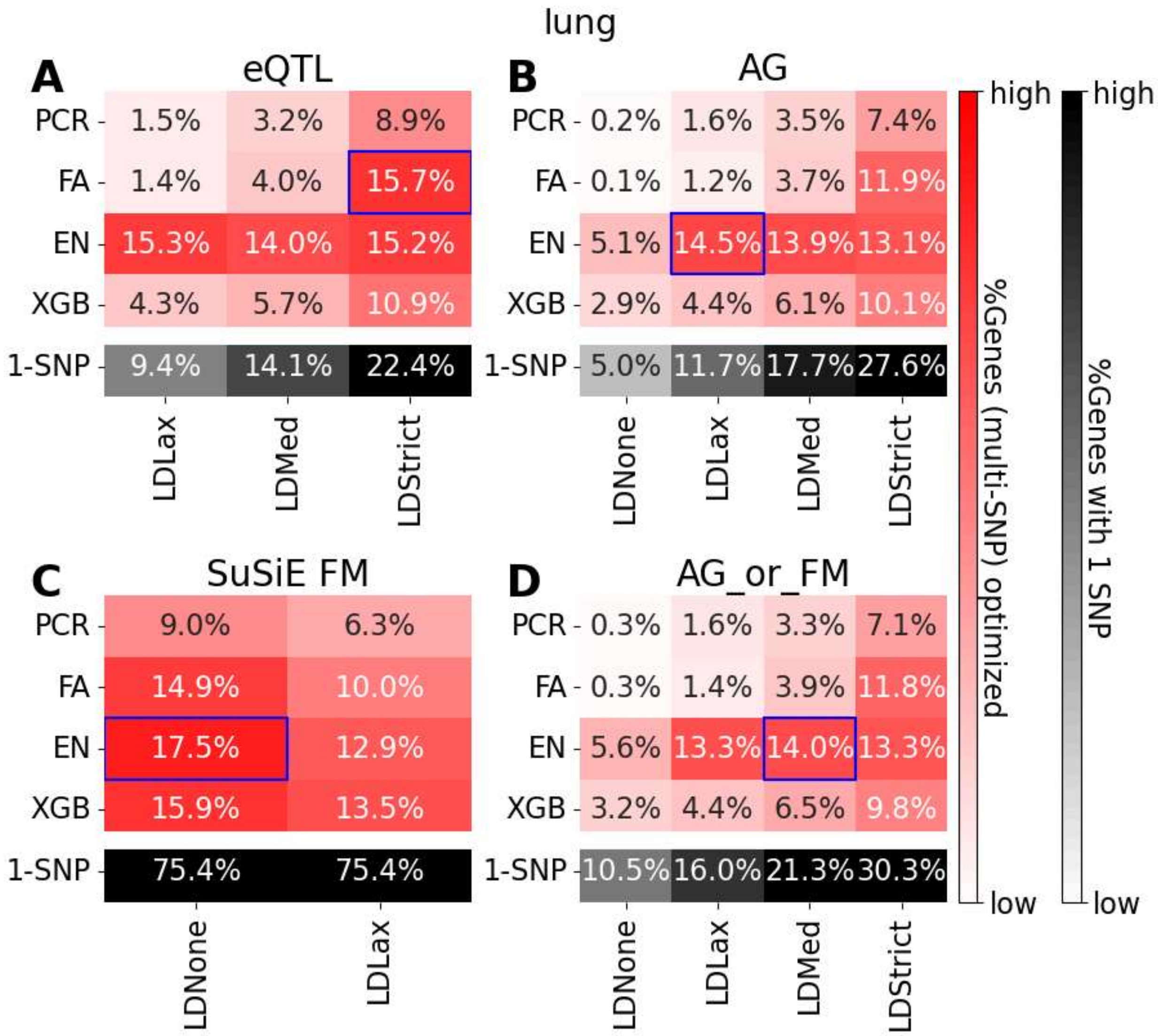


Summary of EAGLES modes impact on adjusted $R^2$ in held-out GTEx lung samples. EAGLES models were trained using GTEx eQTLs (A), AlphaGenome filtered eQTLs (B), GTEx finemapped eQTLs (C) or the union of AlphaGenome and finemapped eQTLs (D). Heatmaps show the fraction of polygenic EAGLES models which maximize performance (adjusted $R^2$ ) relative to other tested EAGLES modes. The black row indicates the fraction of models with a single SNP. Blue boxes indicate the best EAGLES mode. PCR=principal component regression, FA=flip allele count, EN=elasticnet, XGB=xgboost, LD=linkage disequilibrium, FM=finemapped, AG=AlphaGenome.

Figure S5

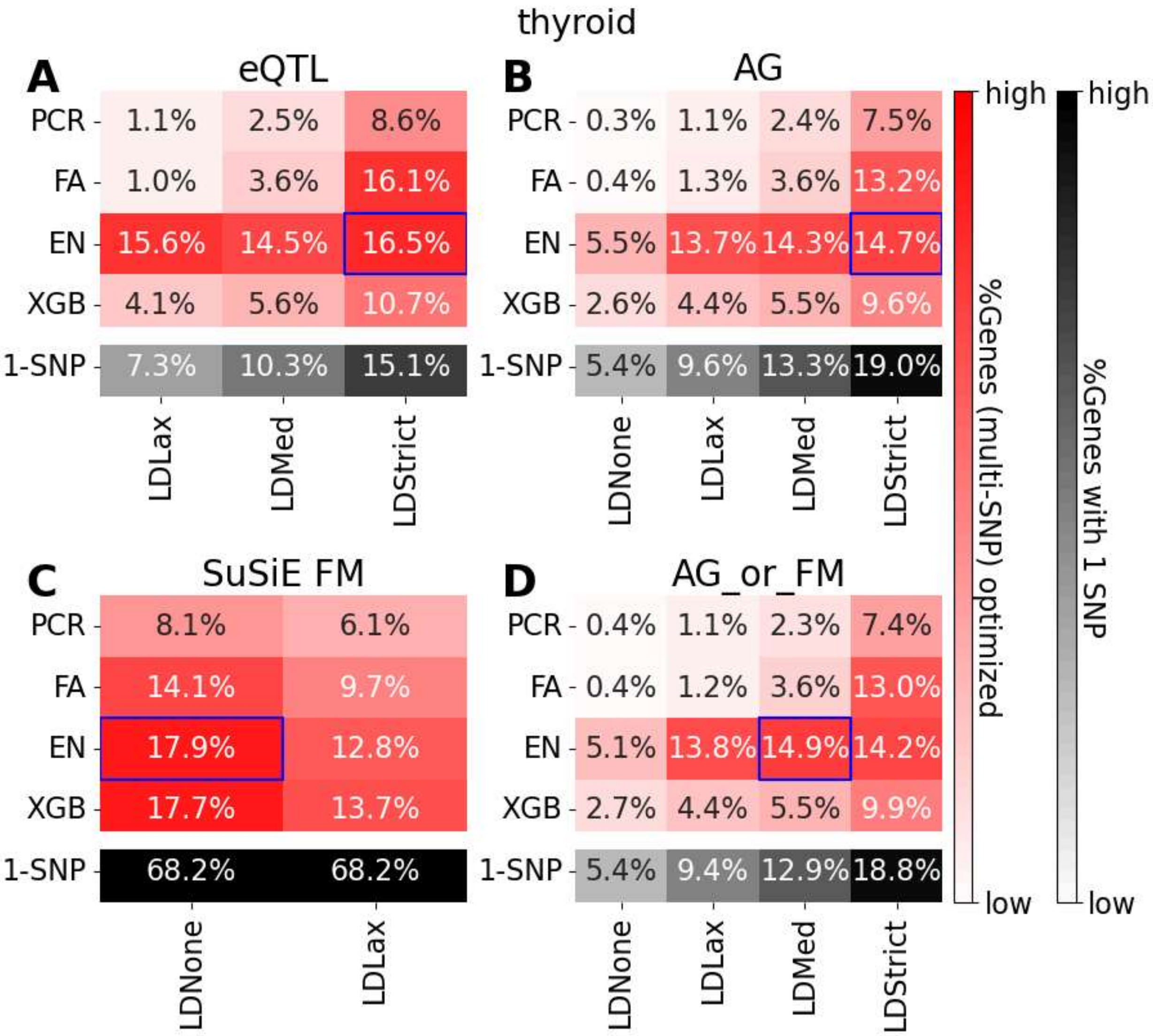


Summary of EAGLES modes impact on adjusted $R^2$ in held-out GTEx thyroid samples. EAGLES models were trained using GTEx eQTLs (A), AlphaGenome filtered eQTLs (B), GTEx finemapped eQTLs (C) or the union of AlphaGenome and finemapped eQTLs (D). Heatmaps show the fraction of polygenic EAGLES models which maximize performance (adjusted $R^2$ ) relative to other tested EAGLES modes. The black row indicates the fraction of models with a single SNP. Blue boxes indicate the best EAGLES mode. PCR=principal component regression, FA=flip allele count, EN=elasticnet, XGB=xgboost, LD=linkage disequilibrium, FM=finemapped, AG=AlphaGenome.

Figure S6

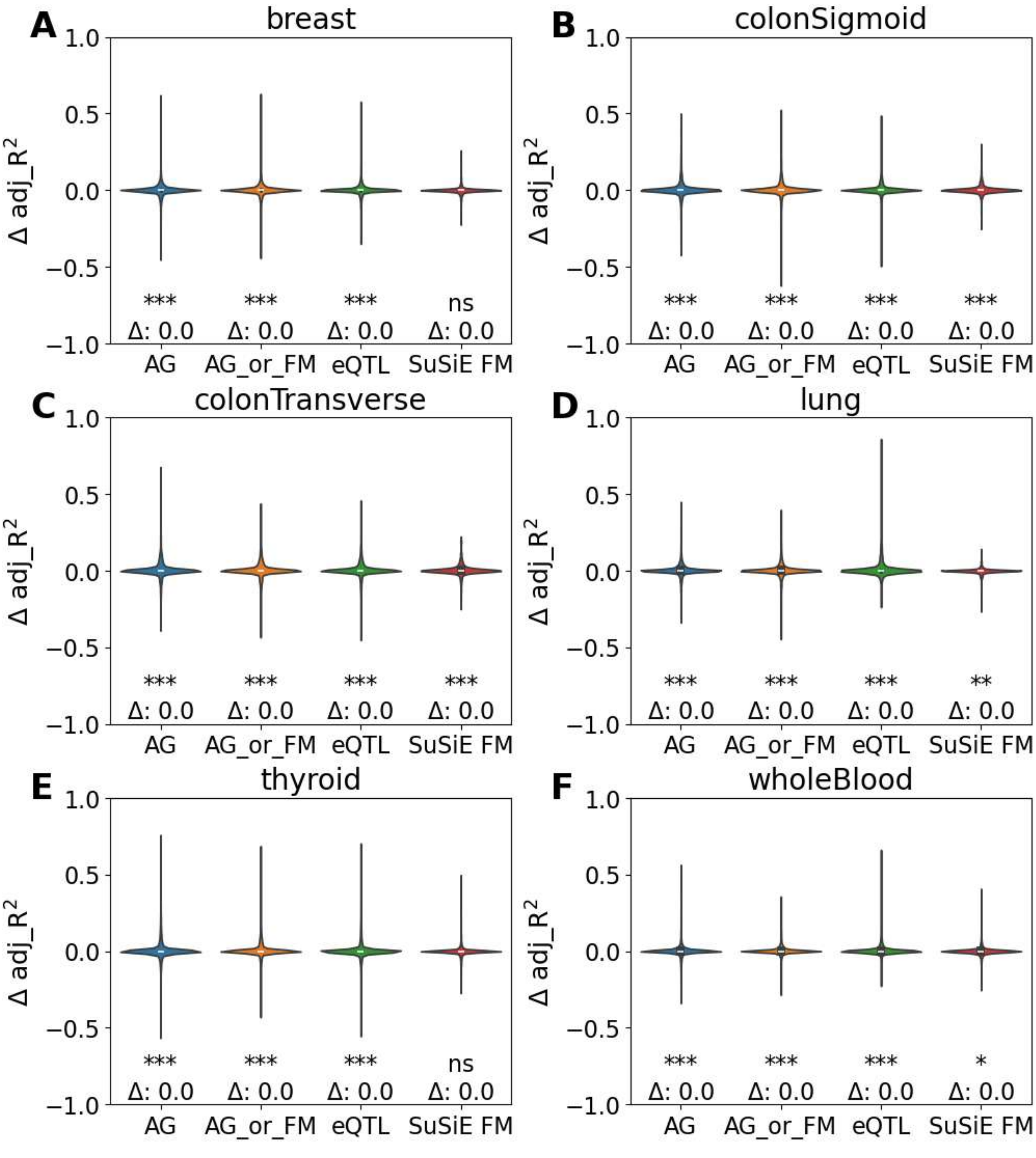


Performance change by choosing EAGLES mode on a per-gene basis. Violin plots summarize the change in EAGLES performance (adjusted $R^2$) in the independent cohorts by selecting the EAGLES mode per-gene rather than a single mode based on held-out GTEx performance. The median performance difference (Δ) is annotated, and Wilcoxon signed-rank tests were used for comparisons: ***<0.001; **<0.01; *<0.05; ns>0.05. AG=AlphaGenome; FM=finemapped.

Figure S7

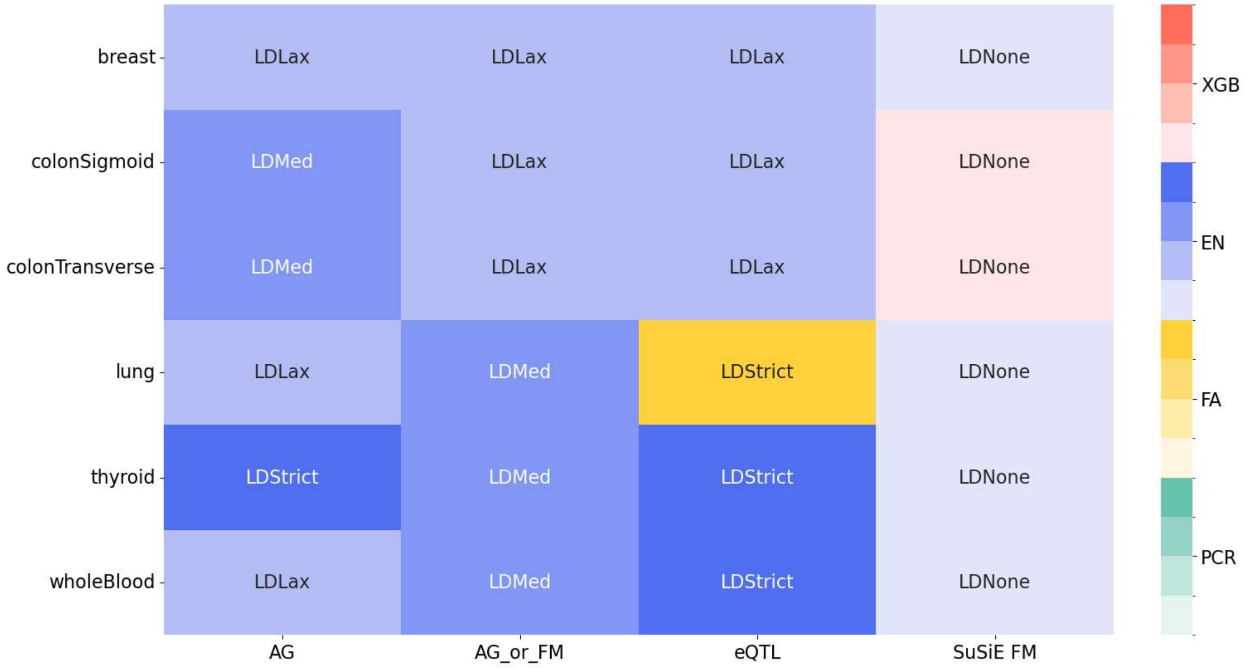


Best EAGLES mode across tissue and eQTL types. Heatmap summarizes the best EAGLES mode (LD pruning threshold and model type) across 6 examined tissue types and 4 eQTL types. The best model type is indicated by color. The best LD pruning threshold is annotated with stricter filtering appearing with a darker shade. FM=finemapped, EN=elasticnet, PCR=principal component regression, XGB=xgboost; AG=AlphaGenome, FA=flip allele count; LD=linkage disequilibrium.

Figure S8

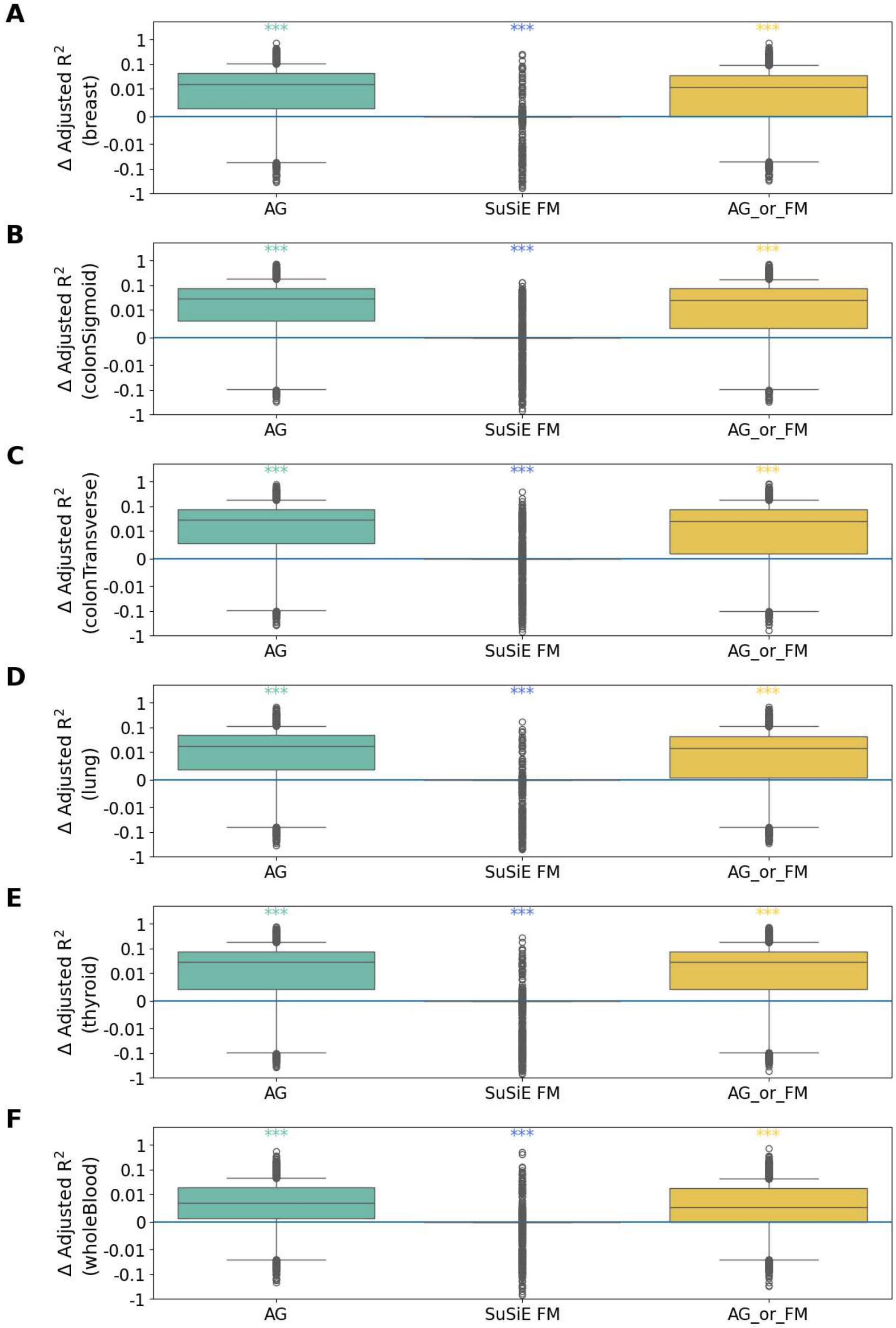


Impact of LD Pruning on EAGLES performance in independent cohorts. For each gene, the difference in EAGLES performance (adjusted $R^2$) in the independent cohorts was computed between the best performing model following LD pruning and the model with no LD pruning. Positive values indicate better performance in the LD pruned model. Wilcoxon signed-rank tests were used for comparisons: ***<0.001; **<0.01; *<0.05; ns>0.05. AG=AlphaGenome; FM=finemapped.

Figure S9

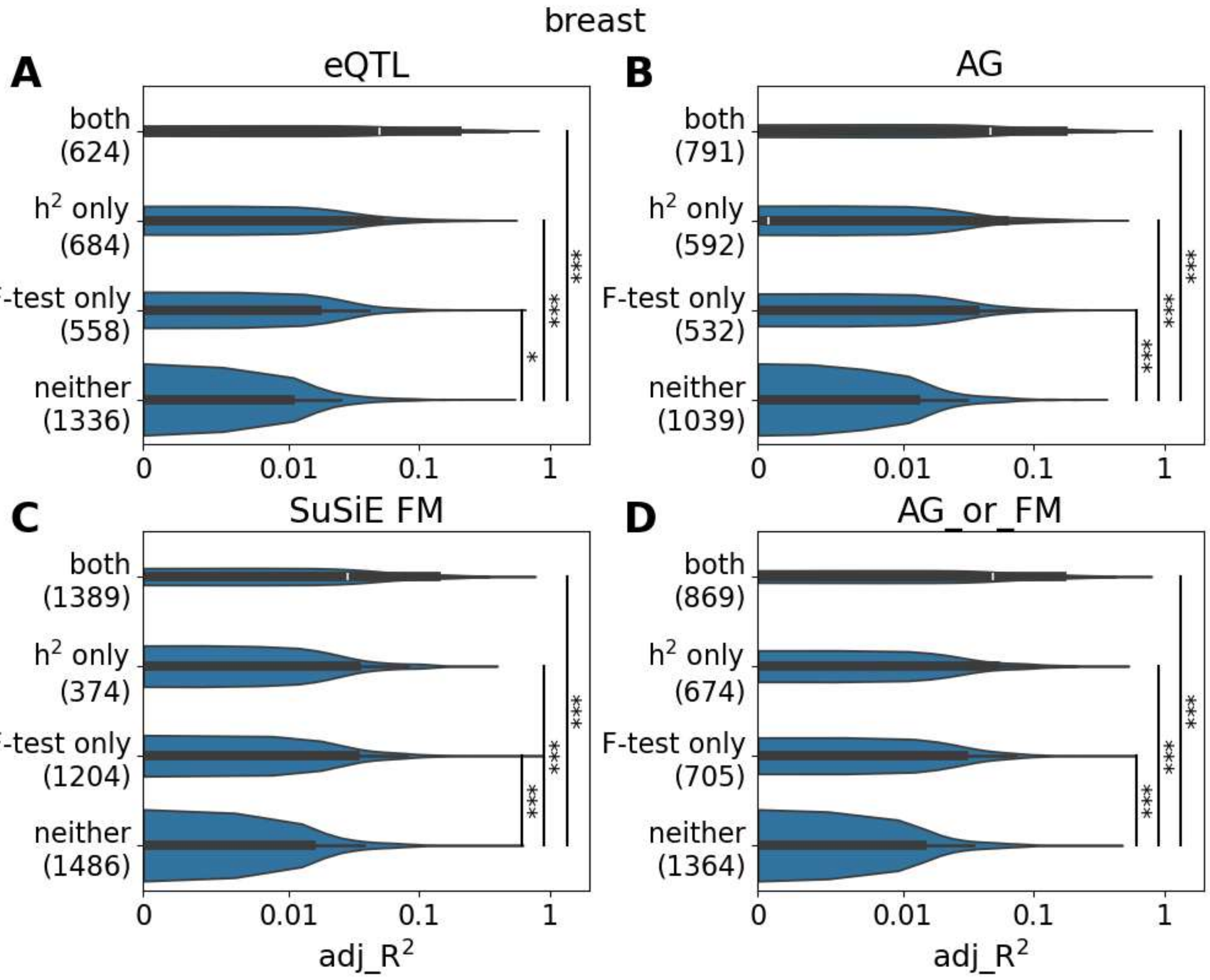


Identifying high-performing EAGLES models in the independent breast cohort. EAGLES models were grouped based on whether they passed the F test and/or predicted expression of genes with significant cis-heritability estimated in GTEx. Models use features from GTEx eQTLs (A), AlphaGenome prioritized eQTLs (B), GTEx finemapped (FM) eQTLs (C) or union of AlphaGenome and finemapped eQTLs (D). Labels on y-axis note F-test and heritability (h2) category with the number of genes per category in parenthesis. Mann-Whitney-U tests (left-tailed) were used for comparisons: ***<0.001; **<0.01; *<0.05; ns>0.05. AG=AlphaGenome; FM=finemapped; h2=significant cis-heritability

Figure S10

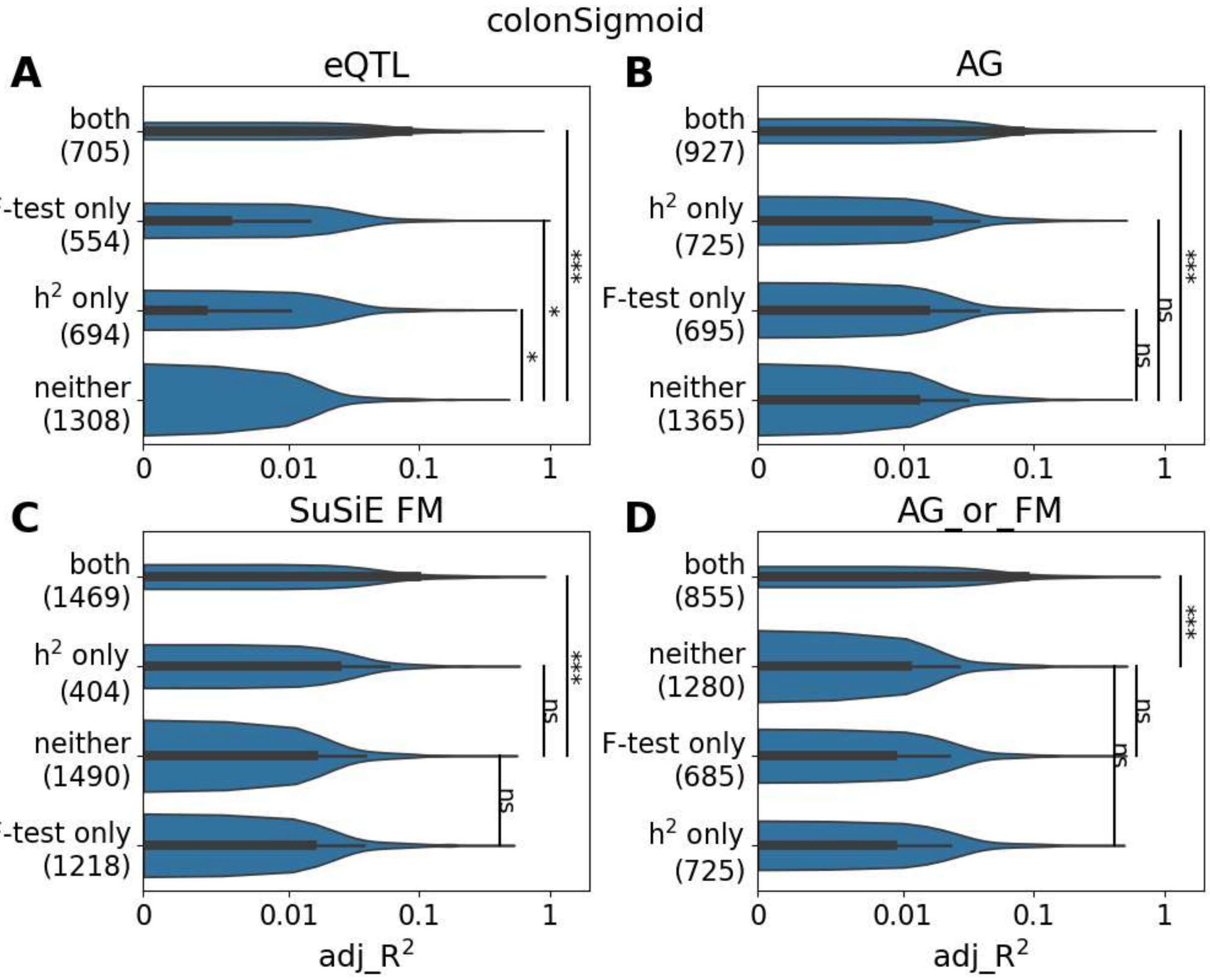


Identifying high-performing sigmoid colon EAGLES models in the independent colon cohort. EAGLES models were grouped based on whether they passed the F test and/or predicted expression of genes with significant cis-heritability estimated in GTEx. Models use features from GTEx eQTLs (A), AlphaGenome prioritized eQTLs (B), GTEx finemapped (FM) eQTLs (C) or union of AlphaGenome and finemapped eQTLs (D). Labels on y-axis note F-test and heritability (h2) category with the number of genes per category in parenthesis. Mann-Whitney-U tests (left-tailed) were used for comparisons: ***<0.001; **<0.01; *<0.05; ns>0.05. AG=AlphaGenome; FM=finemapped; h2=significant cis-heritability

Figure S11

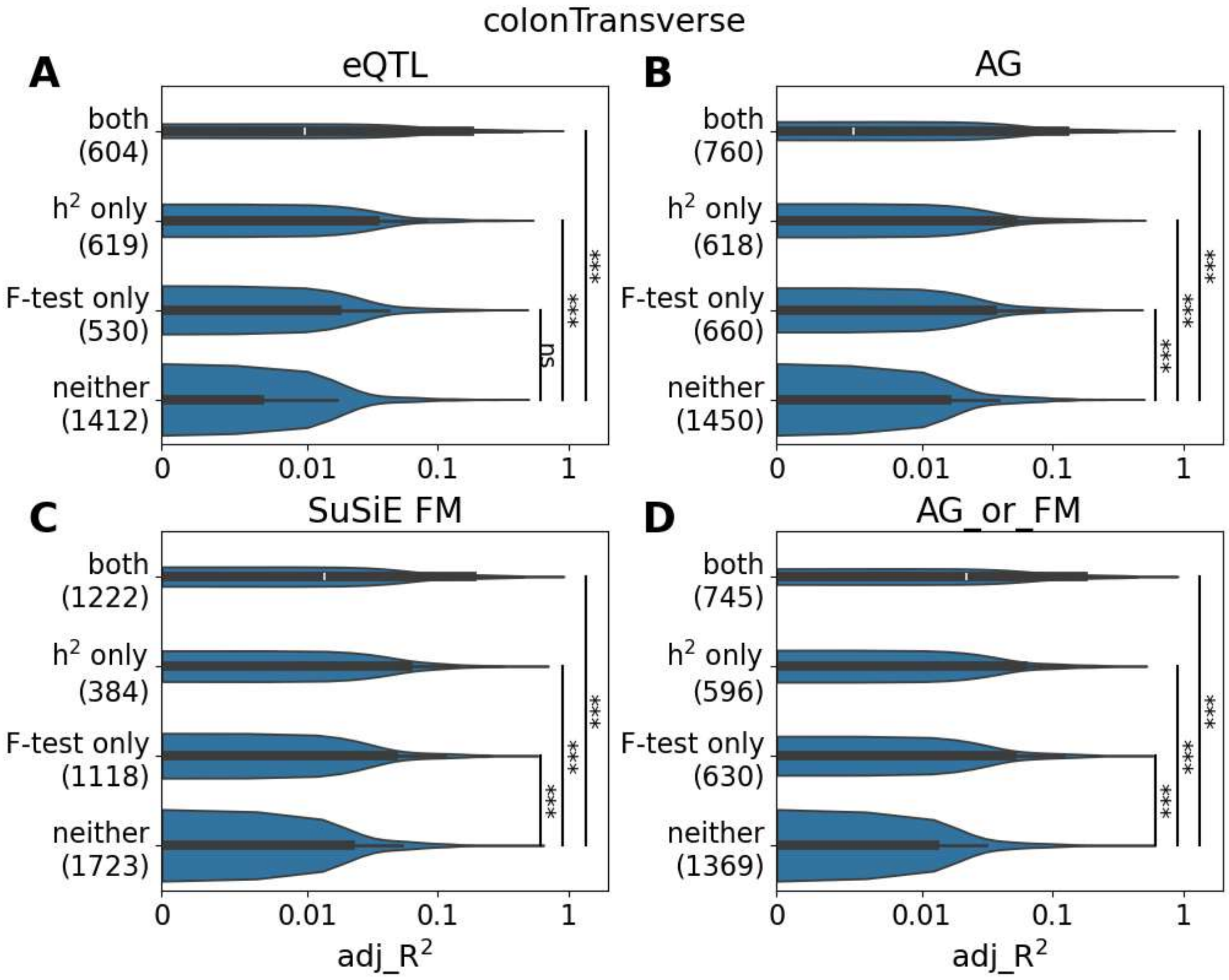


Identifying high-performing transverse colon EAGLES models in the independent colon cohort. EAGLES models were grouped based on whether they passed the F test and/or predicted expression of genes with significant cis-heritability estimated in GTEx. Models use features from GTEx eQTLs (A), AlphaGenome prioritized eQTLs (B), GTEx finemapped (FM) eQTLs (C) or union of AlphaGenome and finemapped eQTLs (D). Labels on y-axis note F-test and heritability (h2) category with the number of genes per category in parenthesis. Mann-Whitney-U tests (left-tailed) were used for comparisons: ***<0.001; **<0.01; *<0.05; ns>0.05. AG=AlphaGenome; FM=finemapped; h2=significant cis-heritability

Figure S12

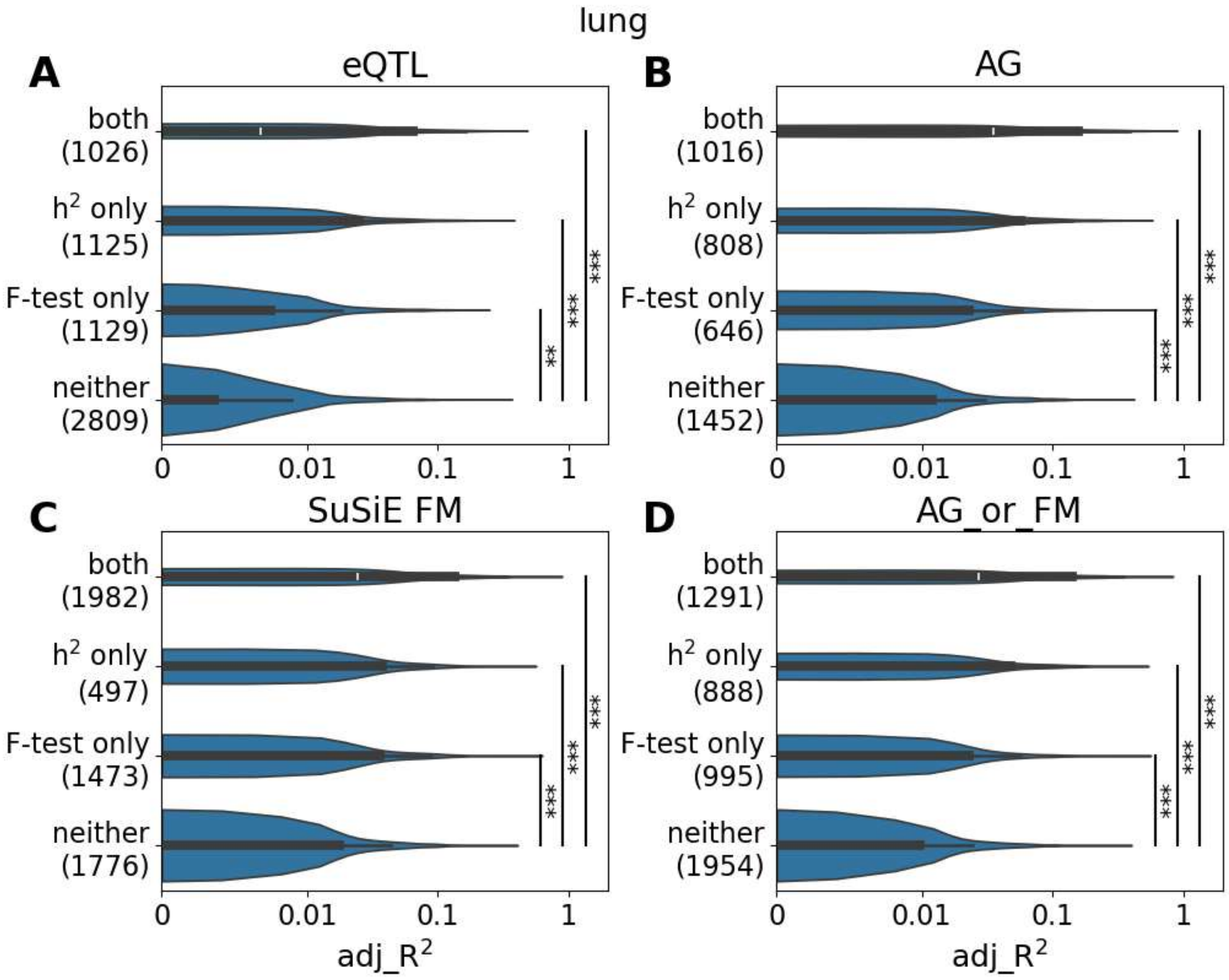


Identifying high-performing EAGLES models in the independent lung cohort. EAGLES models were grouped based on whether they passed the F test and/or predicted expression of genes with significant cis-heritability estimated in GTEx. Models use features from GTEx eQTLs (A), AlphaGenome prioritized eQTLs (B), GTEx finemapped (FM) eQTLs (C) or union of AlphaGenome and finemapped eQTLs (D). Labels on y-axis note F-test and heritability (h2) category with the number of genes per category in parenthesis. Mann-Whitney-U tests (left-tailed) were used for comparisons: ***<0.001; **<0.01; *<0.05; ns>0.05. AG=AlphaGenome; FM=finemapped; h2=significant cis-heritability

Figure S13

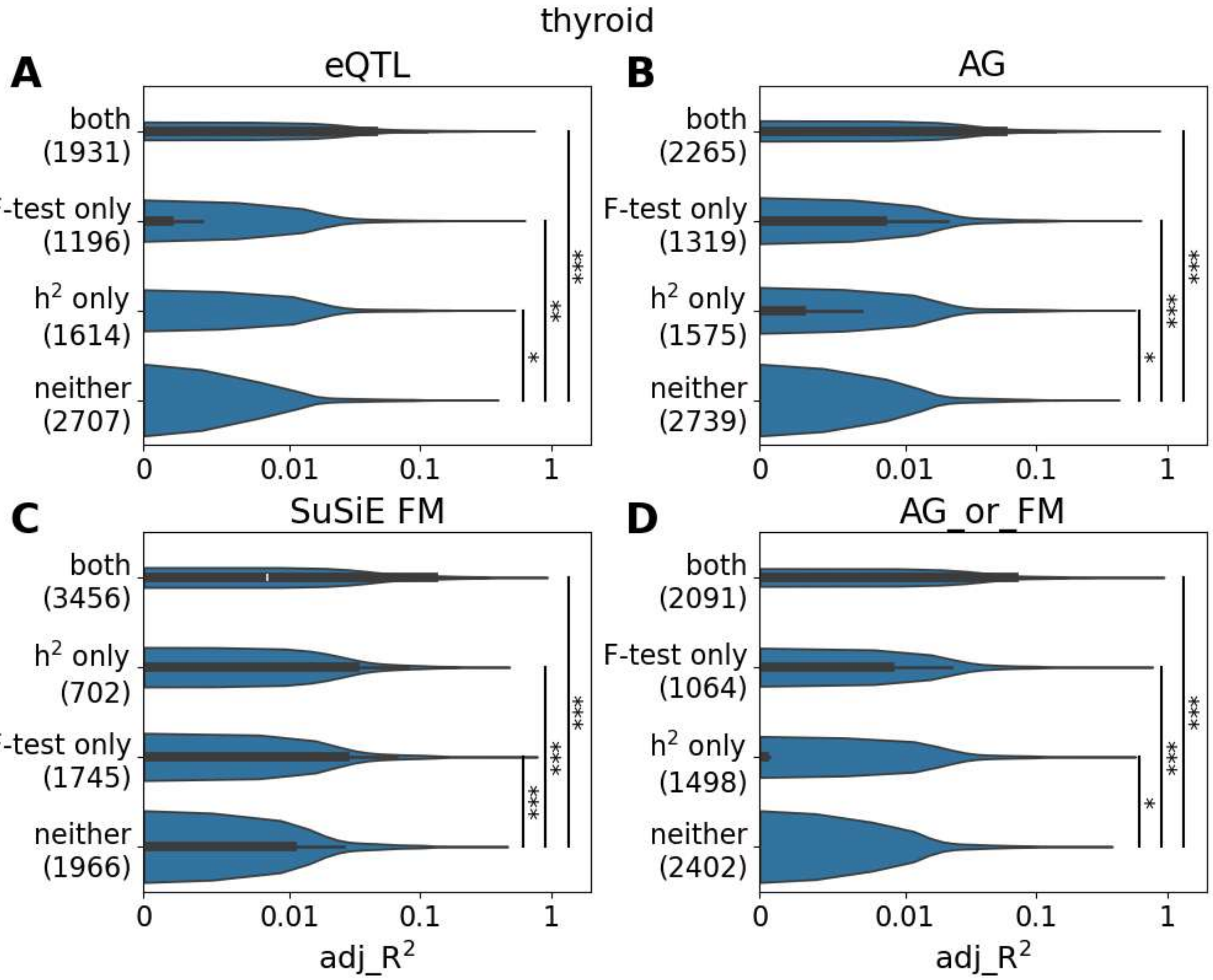


Identifying high-performing EAGLES models in the independent thyroid cohort. EAGLES models were grouped based on whether they passed the F test and/or predicted expression of genes with significant cis-heritability estimated in GTEx. Models use features from GTEx eQTLs (A), AlphaGenome prioritized eQTLs (B), GTEx finemapped (FM) eQTLs (C) or union of AlphaGenome and finemapped eQTLs (D). Labels on y-axis note F-test and heritability (h2) category with the number of genes per category in parenthesis. Mann-Whitney-U tests (left-tailed) were used for comparisons: ***<0.001; **<0.01; *<0.05; ns>0.05. AG=AlphaGenome; FM=finemapped; h2=significant cis-heritability

Figure S14

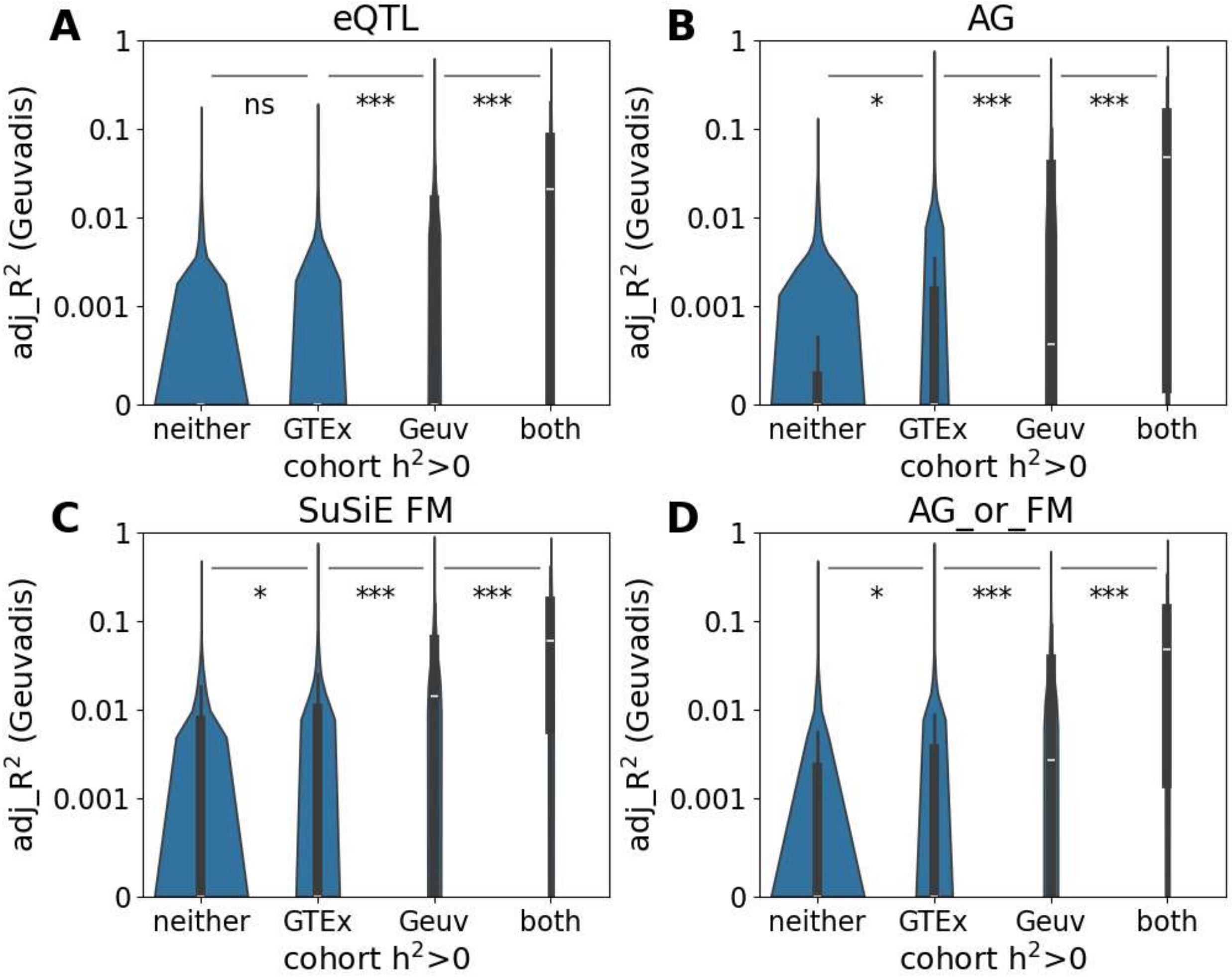


EAGLES model performance and cohort-level heritability differences. Violin plots summarize EAGLES whole blood model performance (adjusted $R^2$) in the independent cohort. Genes were split into four groups based on heritability estimated in each cohort. Mann-Whitney-U tests (left-tailed) were used for comparisons: ***<0.001; **<0.01; *<0.05; ns>0.05. AG=AlphaGenome; FM=finemapped; h2=significant cis-heritability; Geuv=Geuvadis

Figure S15

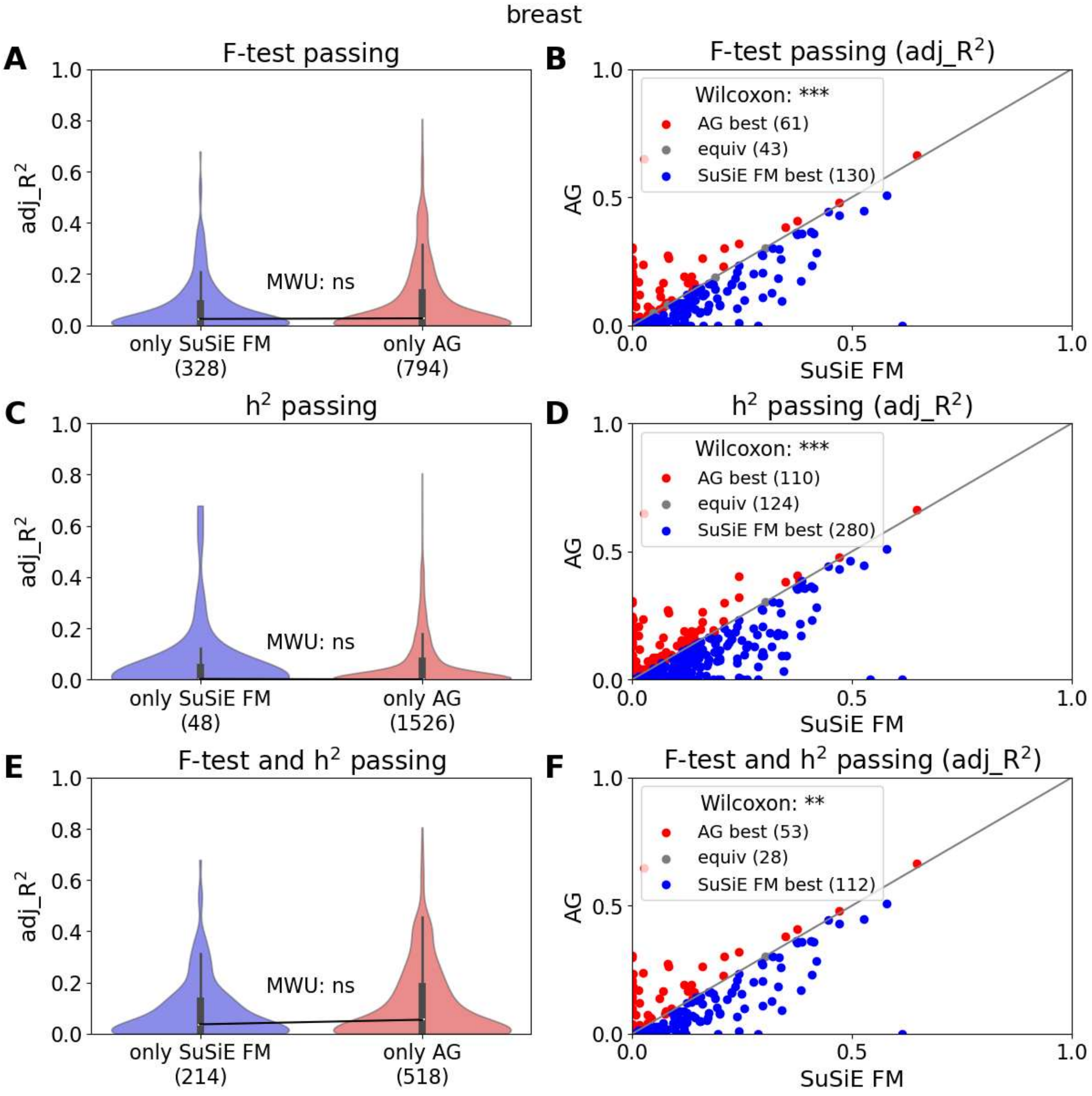


EAGLES breast performance comparison between AG and FM models. Violin plots show EAGLES AG and FM performance (adjusted $R^2$) in the independent breast cohort for models which passed the F-test in held-out GTEx samples (A), showed significant heritability in GTEx (C) or both (E). Violinplots only include genes which satisfy the selection criteria for AG or FM. Mann-Whitney-U (MWU) tests were used for comparisons: ***<0.001; **<0.01; *<0.05; ns>0.05. Scatter plots show EAGLES AG and FM performance for genes satisfying F-test (B), heritability (D) or both (F) criteria for both AG and FM. Wilcoxon signed-rank tests were used for comparisons: ***<0.001; **<0.01; *<0.05; ns>0.05. AG=AlphaGenome; FM=finemapped.

Figure S16

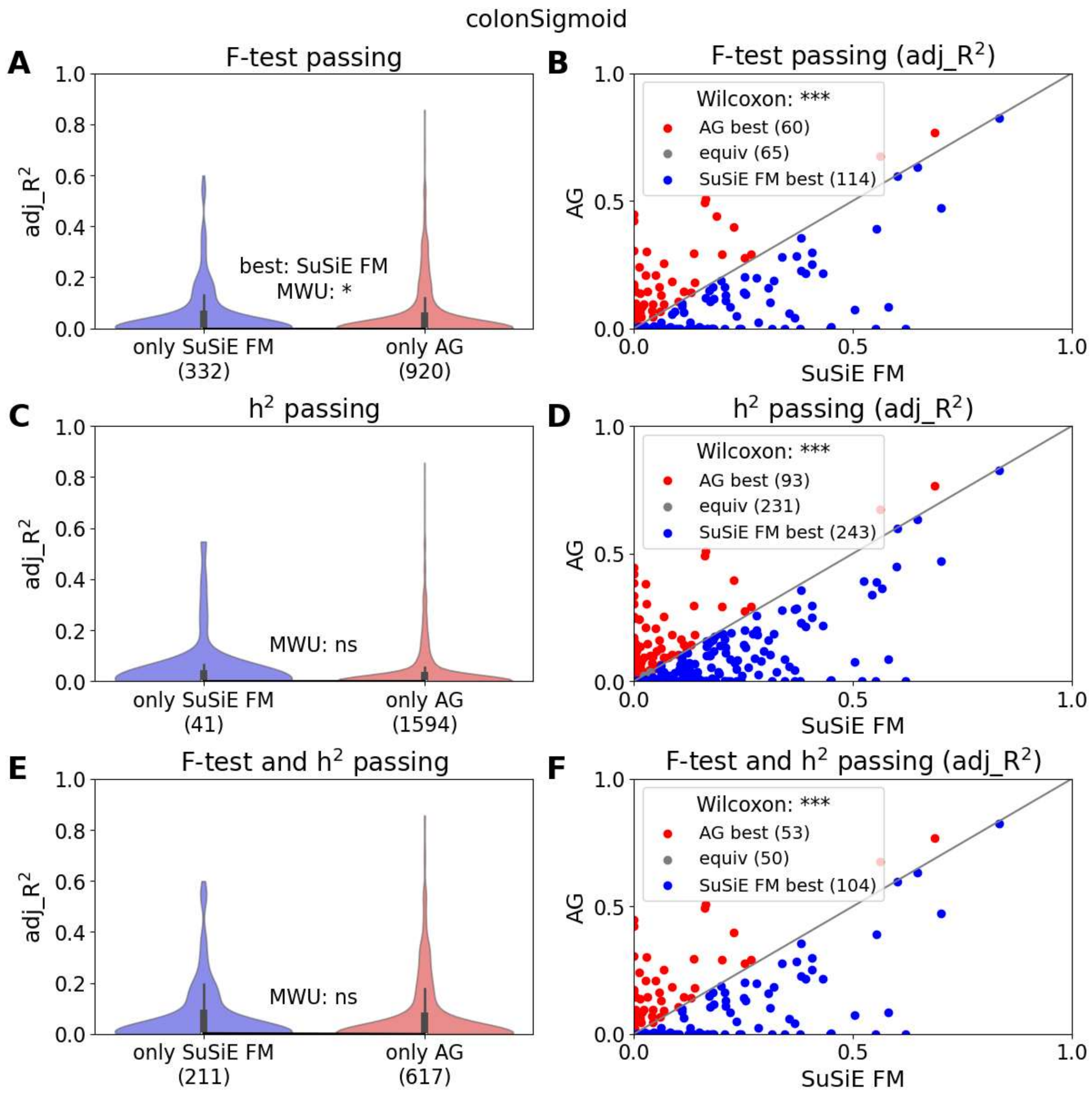


EAGLES breast performance comparison between AG and FM models. Violin plots show EAGLES AG and FM performance (adjusted $R^2$) in the independent colon cohort for models which passed the F-test in held-out GTEx samples (A), showed significant heritability in GTEx (C) or both (E). Violinplots only include genes which satisfy the selection criteria for AG or FM. Mann-Whitney-U (MWU) tests were used for comparisons: ***<0.001; **<0.01; *<0.05; ns>0.05. Scatter plots show EAGLES AG and FM performance for genes satisfying F-test (B), heritability (D) or both (F) criteria for both AG and FM. Wilcoxon signed-rank tests were used for comparisons: ***<0.001; **<0.01; *<0.05; ns>0.05. AG=AlphaGenome; FM=finemapped.

Figure S17

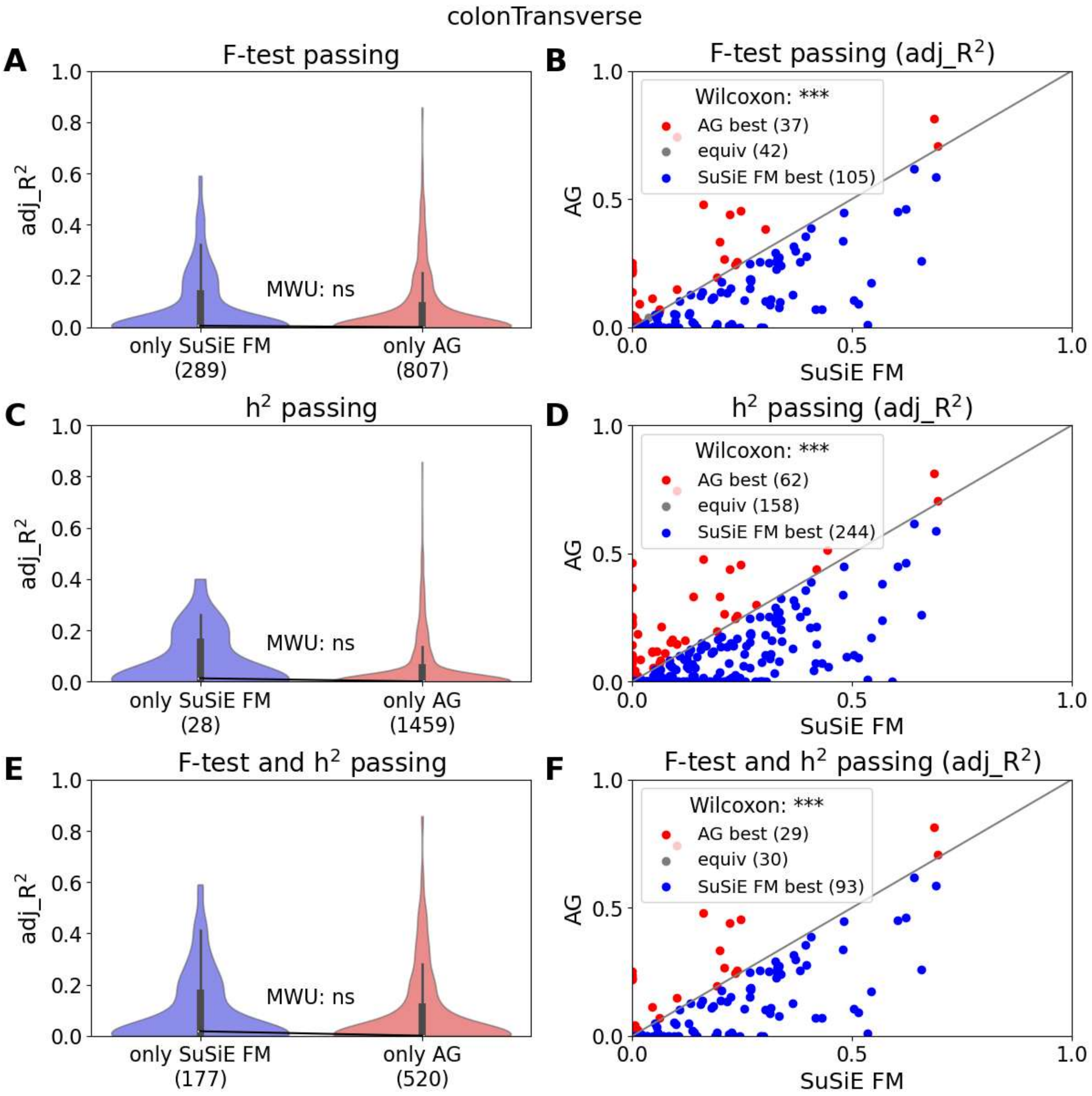


EAGLES breast performance comparison between AG and FM models. Violin plots show EAGLES AG and FM performance (adjusted $R^2$) in the independent colon cohort for models which passed the F-test in held-out GTEx samples (A), showed significant heritability in GTEx (C) or both (E). Violinplots only include genes which satisfy the selection criteria for AG or FM. Mann-Whitney-U (MWU) tests were used for comparisons: ***<0.001; **<0.01; *<0.05; ns>0.05. Scatter plots show EAGLES AG and FM performance for genes satisfying F-test (B), heritability (D) or both (F) criteria for both AG and FM. Wilcoxon signed-rank tests were used for comparisons: ***<0.001; **<0.01; *<0.05; ns>0.05. AG=AlphaGenome; FM=finemapped.

Figure S18

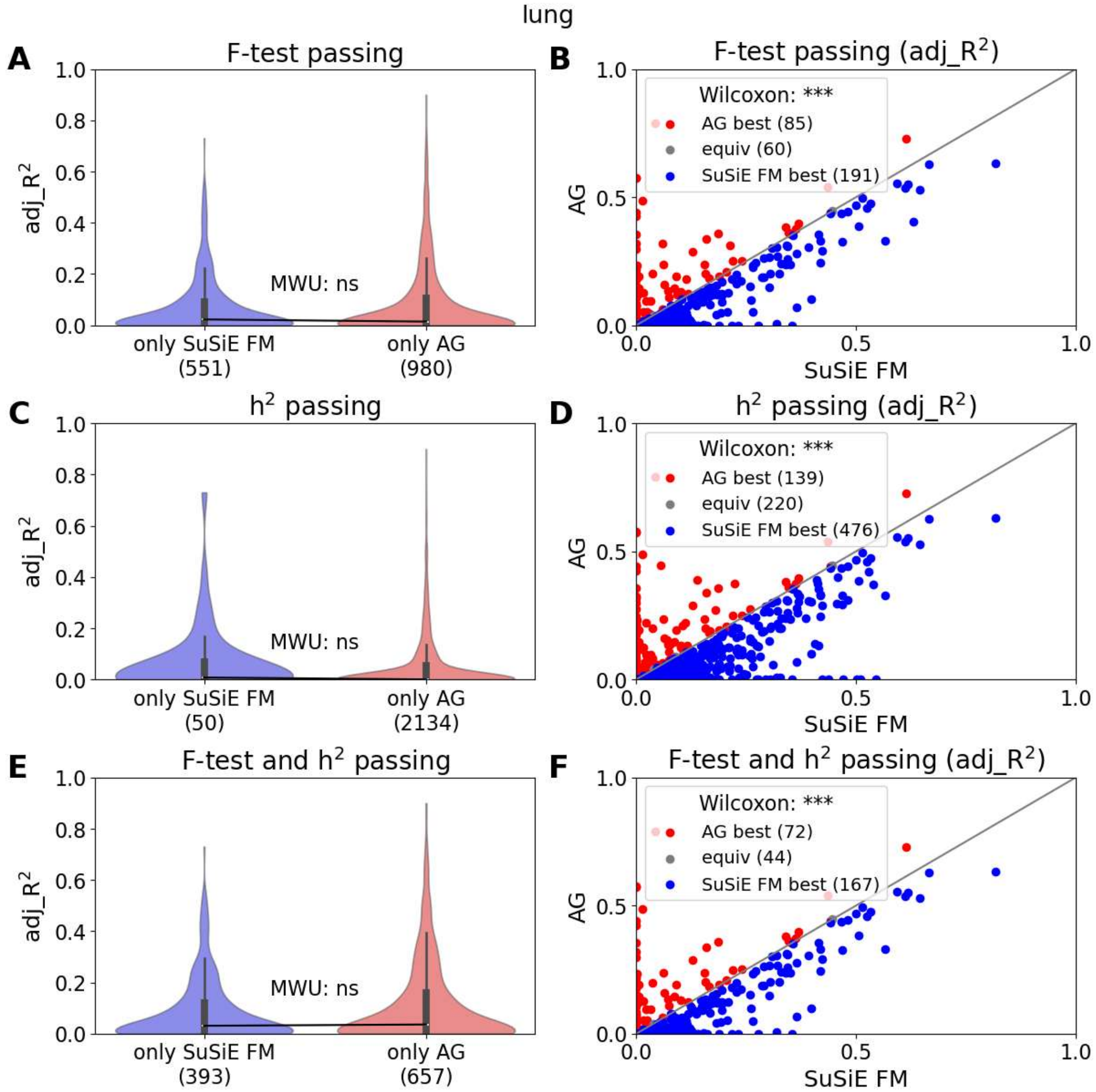


EAGLES breast performance comparison between AG and FM models. Violin plots show EAGLES AG and FM performance (adjusted $R^2$) in the independent lung cohort for models which passed the F-test in held-out GTEx samples (A), showed significant heritability in GTEx (C) or both (E). Violinplots only include genes which satisfy the selection criteria for AG or FM. Mann-Whitney-U (MWU) tests were used for comparisons: ***<0.001; **<0.01; *<0.05; ns>0.05. Scatter plots show EAGLES AG and FM performance for genes satisfying F-test (B), heritability (D) or both (F) criteria for both AG and FM. Wilcoxon signed-rank tests were used for comparisons: ***<0.001; **<0.01; *<0.05; ns>0.05. AG=AlphaGenome; FM=finemapped.

Figure S19

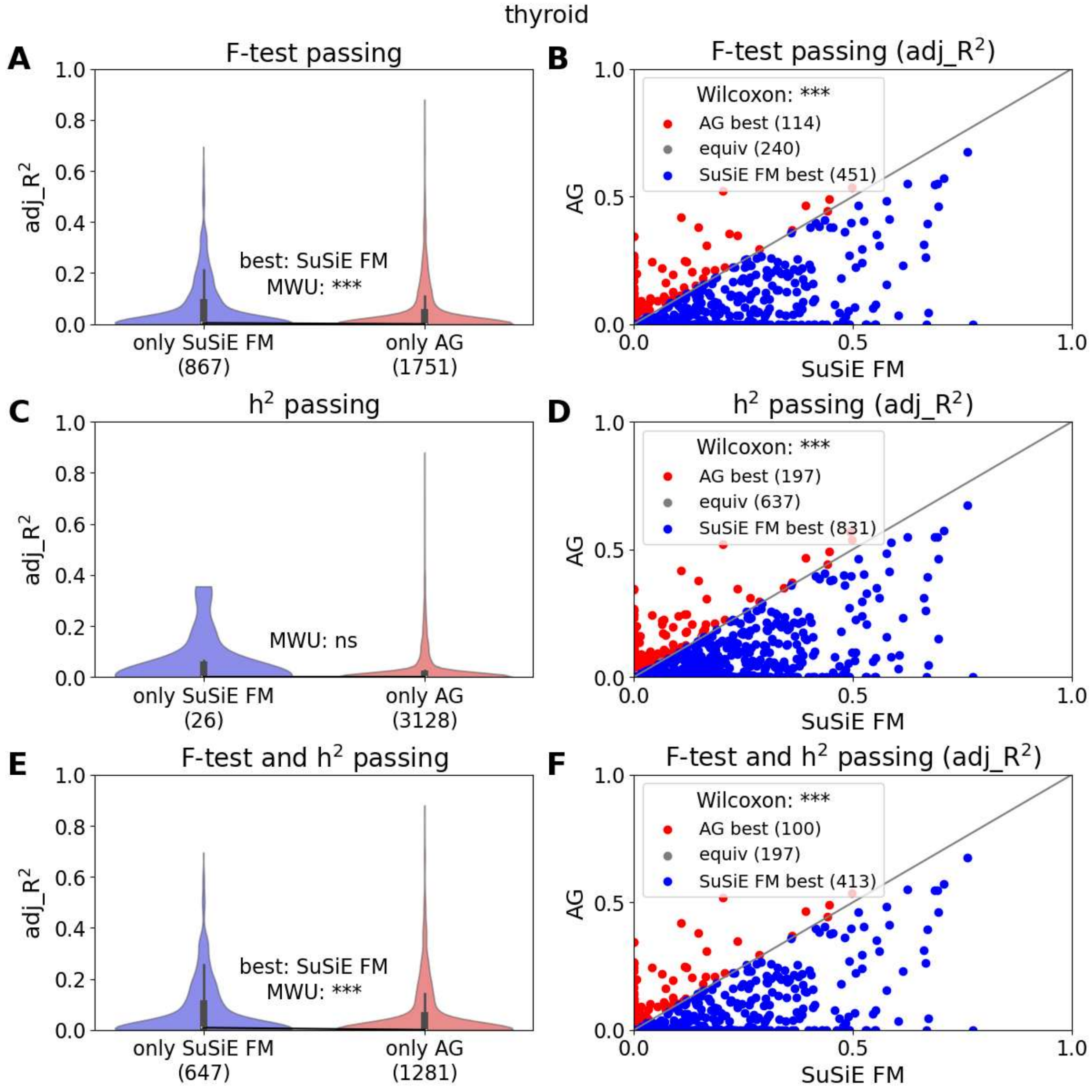


EAGLES breast performance comparison between AG and FM models. Violin plots show EAGLES AG and FM performance (adjusted $R^2$) in the independent thyroid cohort for models which passed the F-test in held-out GTEx samples (A), showed significant heritability in GTEx (C) or both (E). Violinplots only include genes which satisfy the selection criteria for AG or FM. Mann-Whitney-U (MWU) tests were used for comparisons: ***<0.001; **<0.01; *<0.05; ns>0.05. Scatter plots show EAGLES AG and FM performance for genes satisfying F-test (B), heritability (D) or both (F) criteria for both AG and FM. Wilcoxon signed-rank tests were used for comparisons: ***<0.001; **<0.01; *<0.05; ns>0.05. AG=AlphaGenome; FM=finemapped.

Figure S20

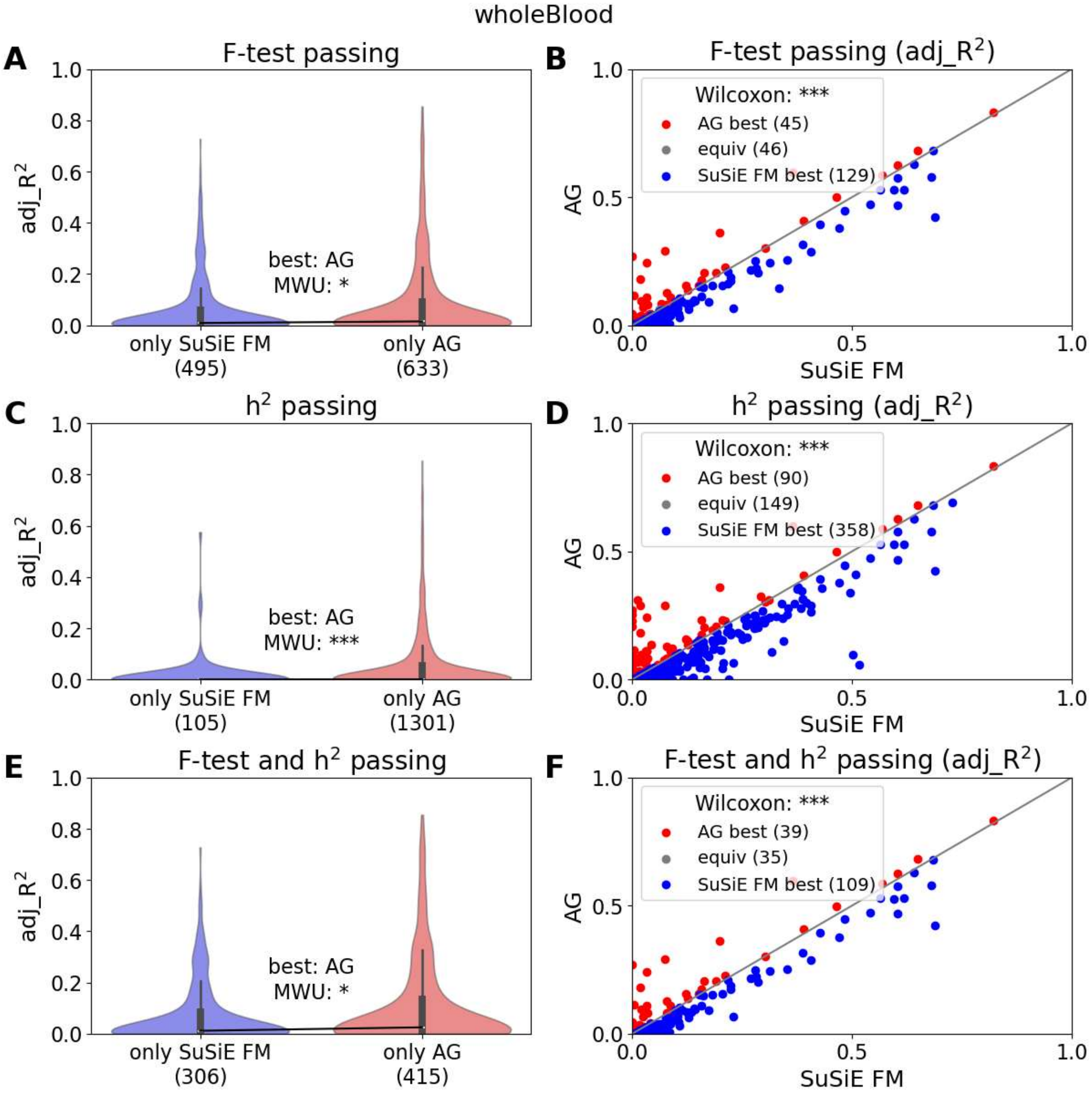


EAGLES breast performance comparison between AG and FM models. Violin plots show EAGLES AG and FM performance (adjusted $R^2$) in the independent blood cohort for models which passed the F-test in held-out GTEx samples (A), showed significant heritability in GTEx (C) or both (E). Violinplots only include genes which satisfy the selection criteria for AG or FM. Mann-Whitney-U (MWU) tests were used for comparisons: ***<0.001; **<0.01; *<0.05; ns>0.05. Scatter plots show EAGLES AG and FM performance for genes satisfying F-test (B), heritability (D) or both (F) criteria for both AG and FM. Wilcoxon signed-rank tests were used for comparisons: ***<0.001; **<0.01; *<0.05; ns>0.05. AG=AlphaGenome; FM=finemapped.

Figure S21

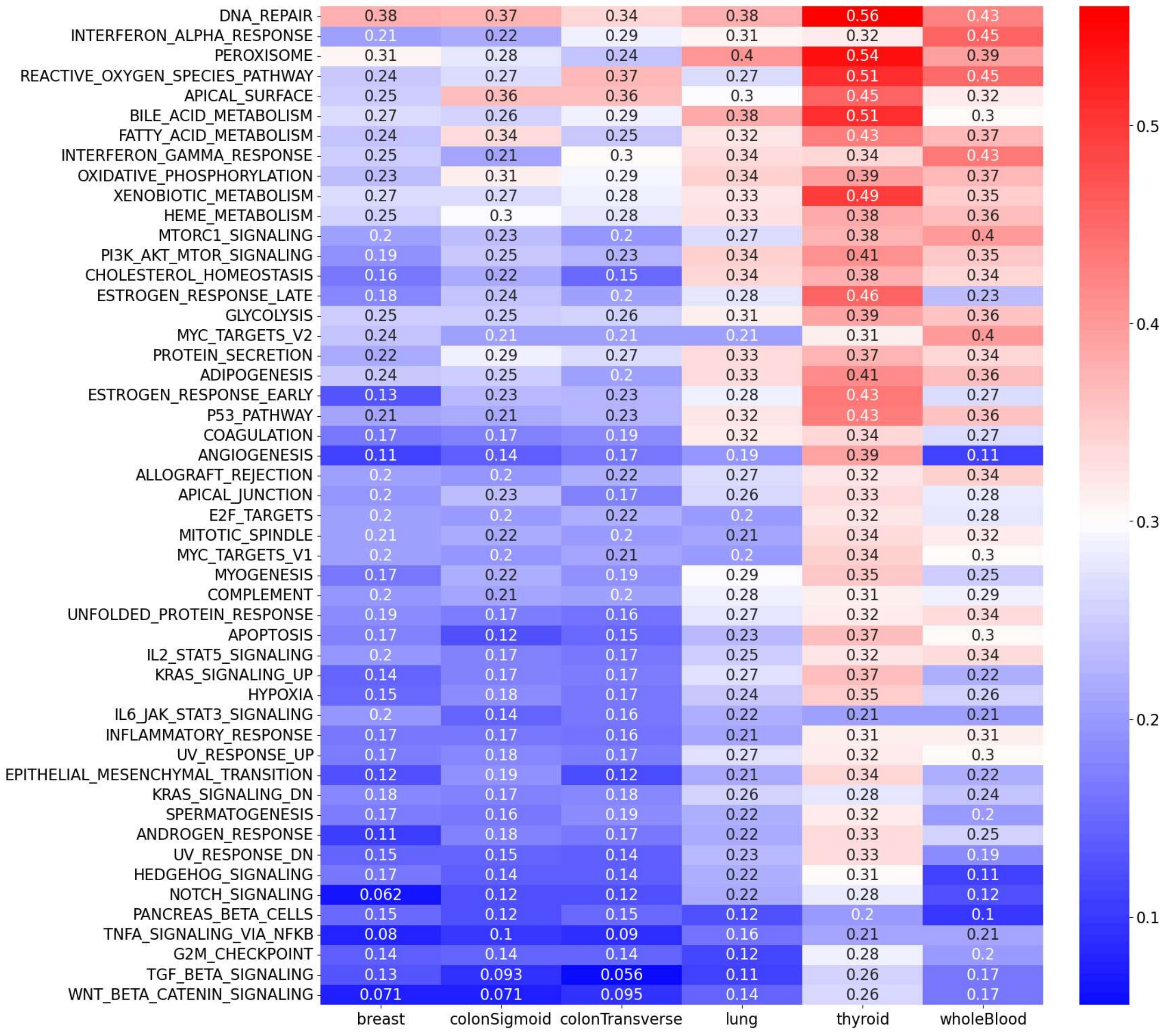

Fraction of MSigDB Hallmark gene sets available in EAGLES. For each tissue and gene set, the lowest proportion of gene set members available in EAGLES was determined across the four examined eQTL sources. These proportions are summarized in this heatmap, with colors chosen to emphasize gene sets that are relatively well represented in EAGLES (red) and not well represented (blue).

Figure S22

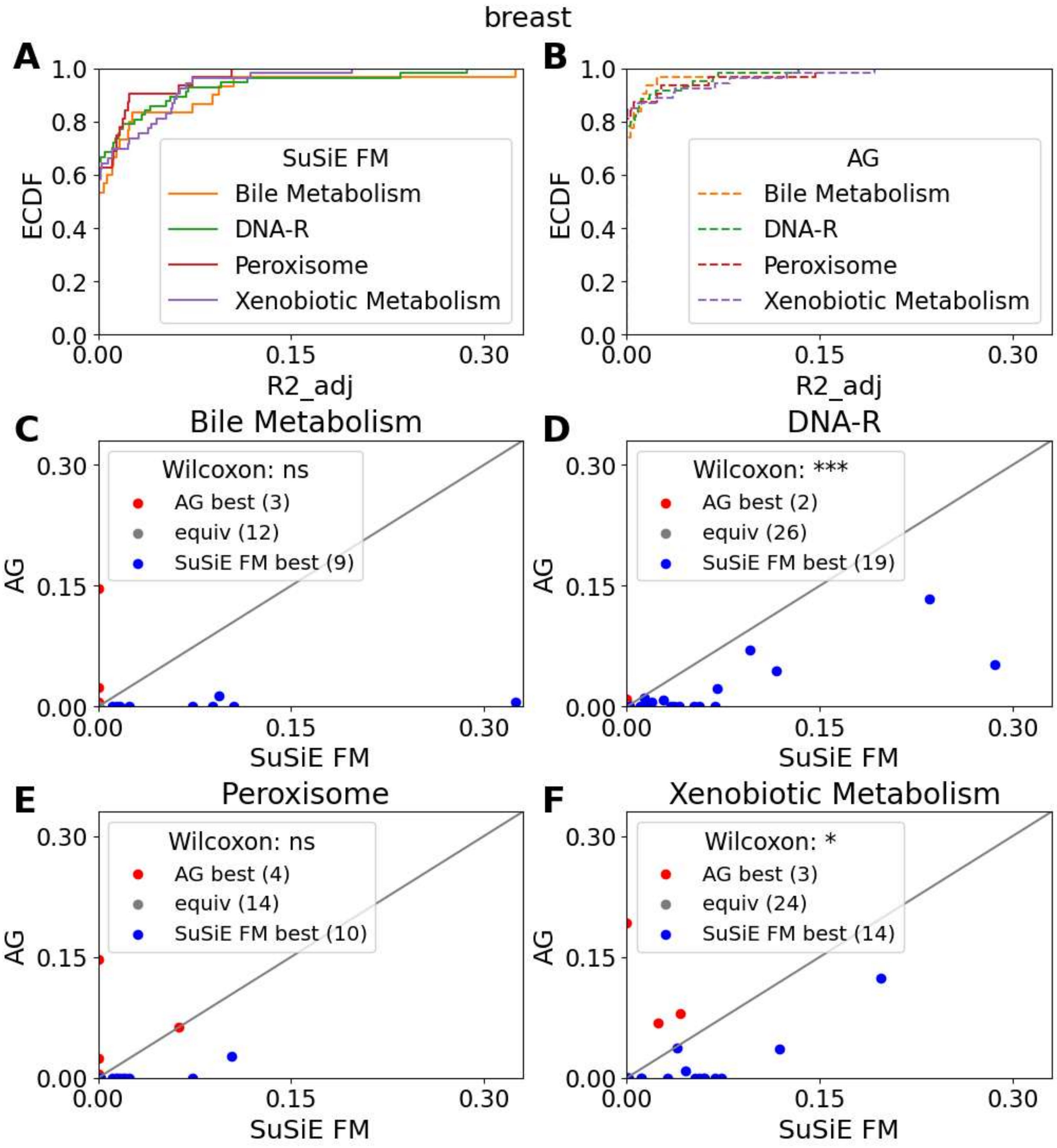


EAGLES coverage in MSigDB Hallmark gene sets. ECDF curves show EAGLES FM (A) and AG (B) performance (adjusted $R^2$) in the independent breast cohort over the top 4 gene sets by fraction of members included in EAGLES. For genes modelable in both EAGLES FM and EAGLES AG, scatter plots compare EAGLES FM and AG performance in bile metabolism (C), DNA repair (D), Peroxisome (E), and Xenobiotic Metabolism (F) genes. The Wilcoxon signed-rank test was used for comparisons: ***<0.001; **<0.01; *<0.05; ns>0.05. FM=finemapped; AG=AlphaGenome; DNA-R=DNA Repair.

Figure S23

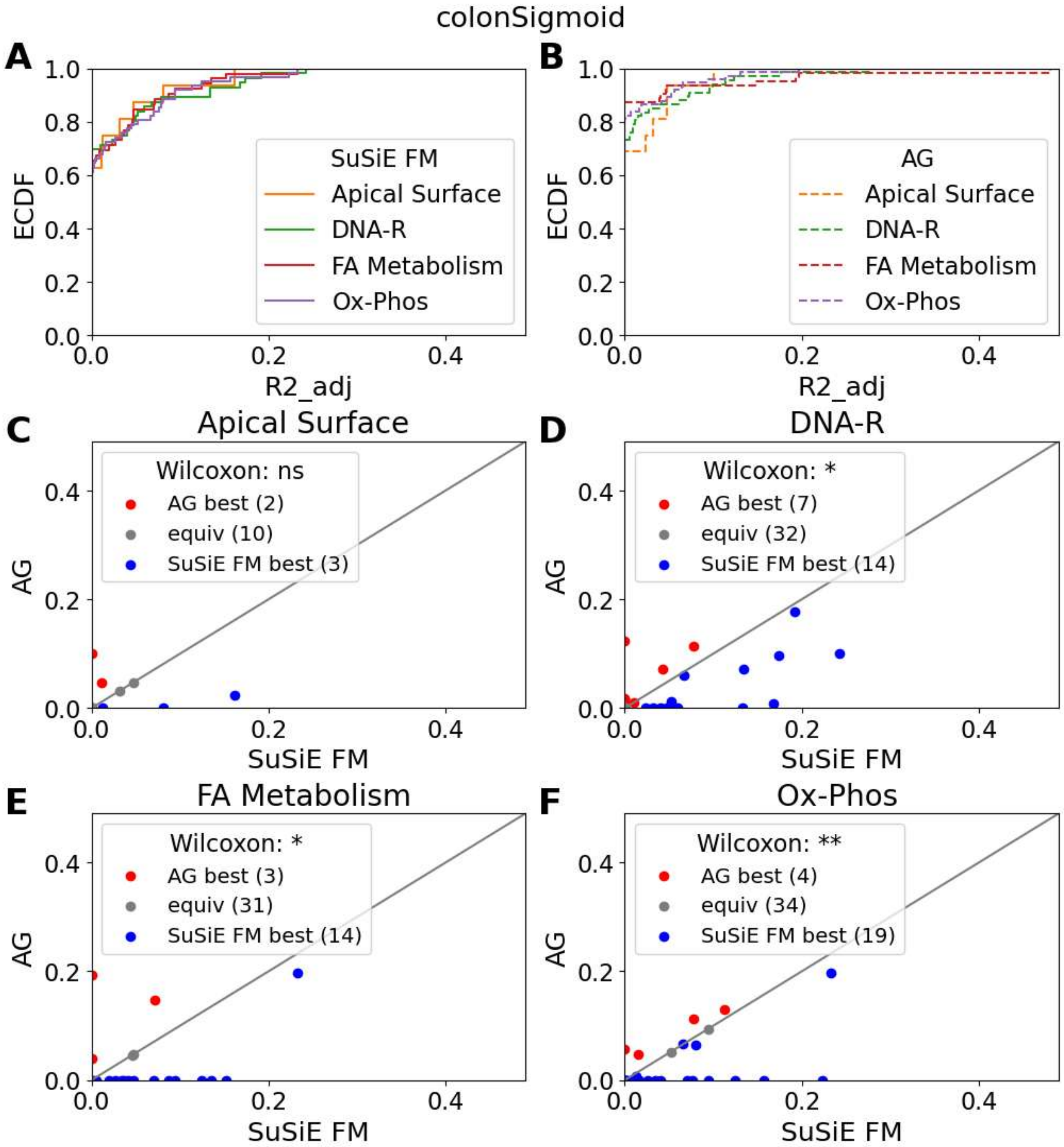


EAGLES coverage in MSigDB Hallmark gene sets. ECDF curves show EAGLES FM (A) and AG (B) performance (adjusted $R^2$) in the independent colon cohort over the top 4 gene sets by fraction of members included in EAGLES. For genes modelable in both EAGLES FM and EAGLES AG, scatter plots compare performance in apical surface (C), DNA repair (D), fatty acid metabolism (E), and oxidative phosphorylation (F) genes. The Wilcoxon signed-rank test was used for comparisons: ***<0.001; **<0.01; *<0.05; ns>0.05. FM=finemapped; AG=AlphaGenome; DNA-R=DNA Repair; FA=fatty acid; Ox-Phos=Oxidative Phosphorylation.

Figure S24

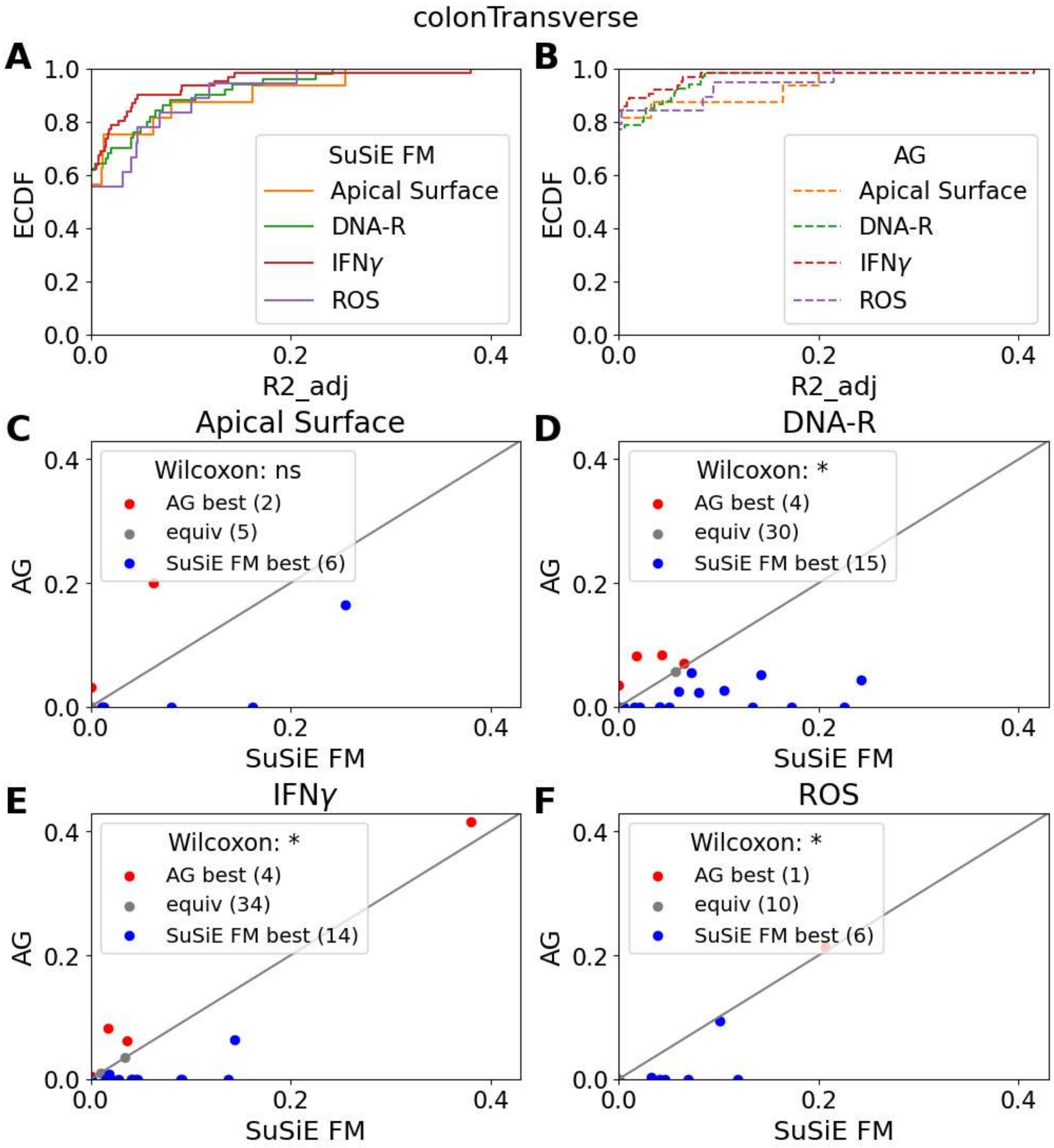


EAGLES coverage in MSigDB Hallmark gene sets. ECDF curves show EAGLES FM (A) and AG (B) performance (adjusted $R^2$) in the independent colon cohort over the top 4 gene sets by fraction of members included in EAGLES. For genes modelable in both EAGLES FM and EAGLES AG, scatter plots compare performance in apical surface (C), DNA repair (D), interferon-$\gamma$ (E), and reactive oxygen species (F) genes. The Wilcoxon signed-rank test was used for comparisons: ***<0.001; **<0.01; *<0.05; ns>0.05. FM=finemapped; AG=AlphaGenome; DNA-R=DNA repair; IFN$\gamma$=interferon-$\gamma$; ROS=reactive oxygen species.

Figure S25

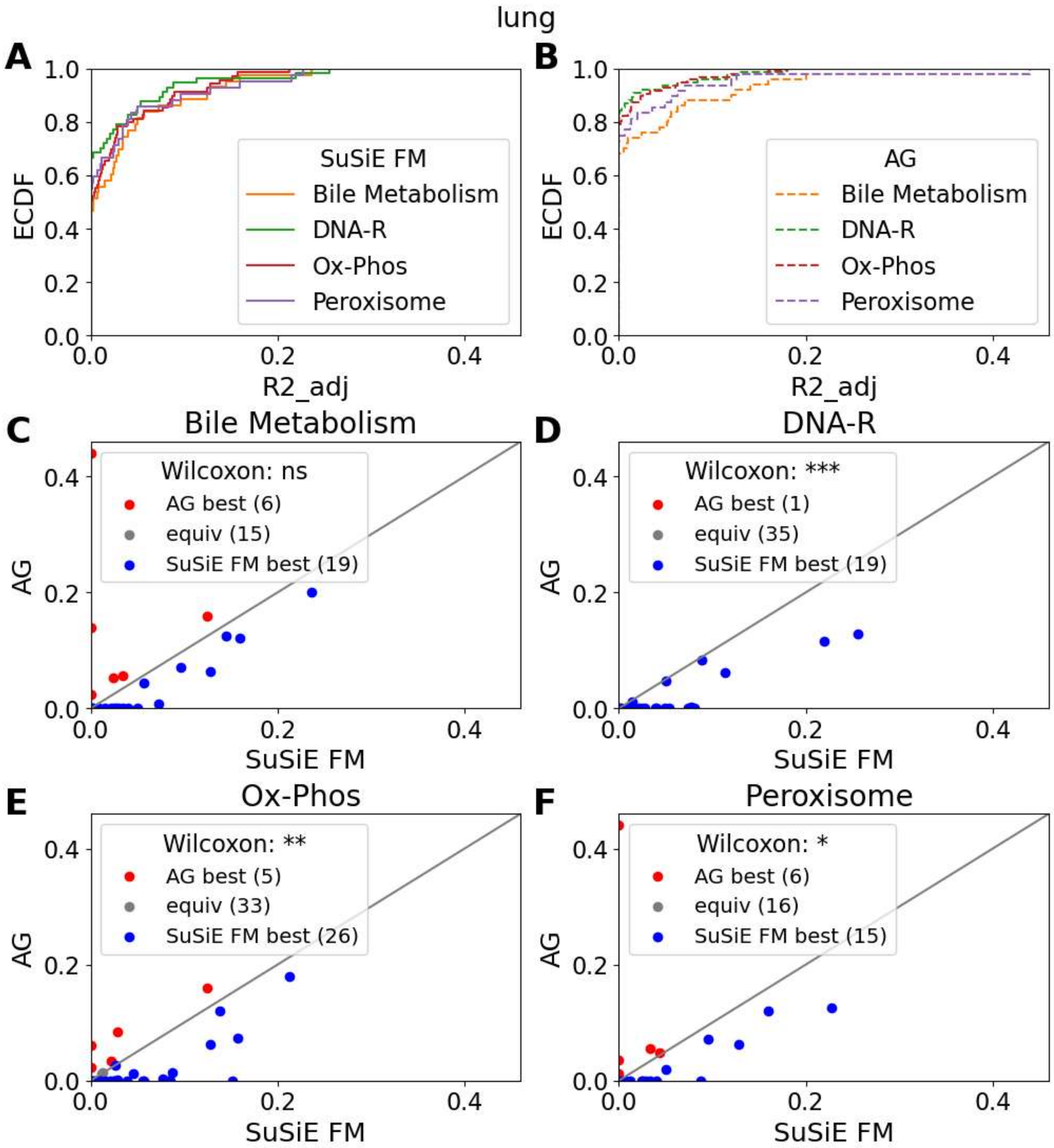


EAGLES coverage in MSigDB Hallmark gene sets. ECDF curves show EAGLES FM (A) and AG (B) performance (adjusted $R^2$) in the independent lung cohort over the top 4 gene sets by fraction of members included in EAGLES. For genes modelable in both EAGLES FM and EAGLES AG, scatter plots compare performance in bile metabolism (C), DNA repair (D), oxidative phosphorylation (E), and peroxisome (F) genes. The Wilcoxon signed-rank test was used for comparisons: ***<0.001; **<0.01; *<0.05; ns>0.05. FM=finemapped; AG=AlphaGenome; DNA-R=DNA repair; Ox-Phos=Oxidative Phosphorylation.

Figure S26

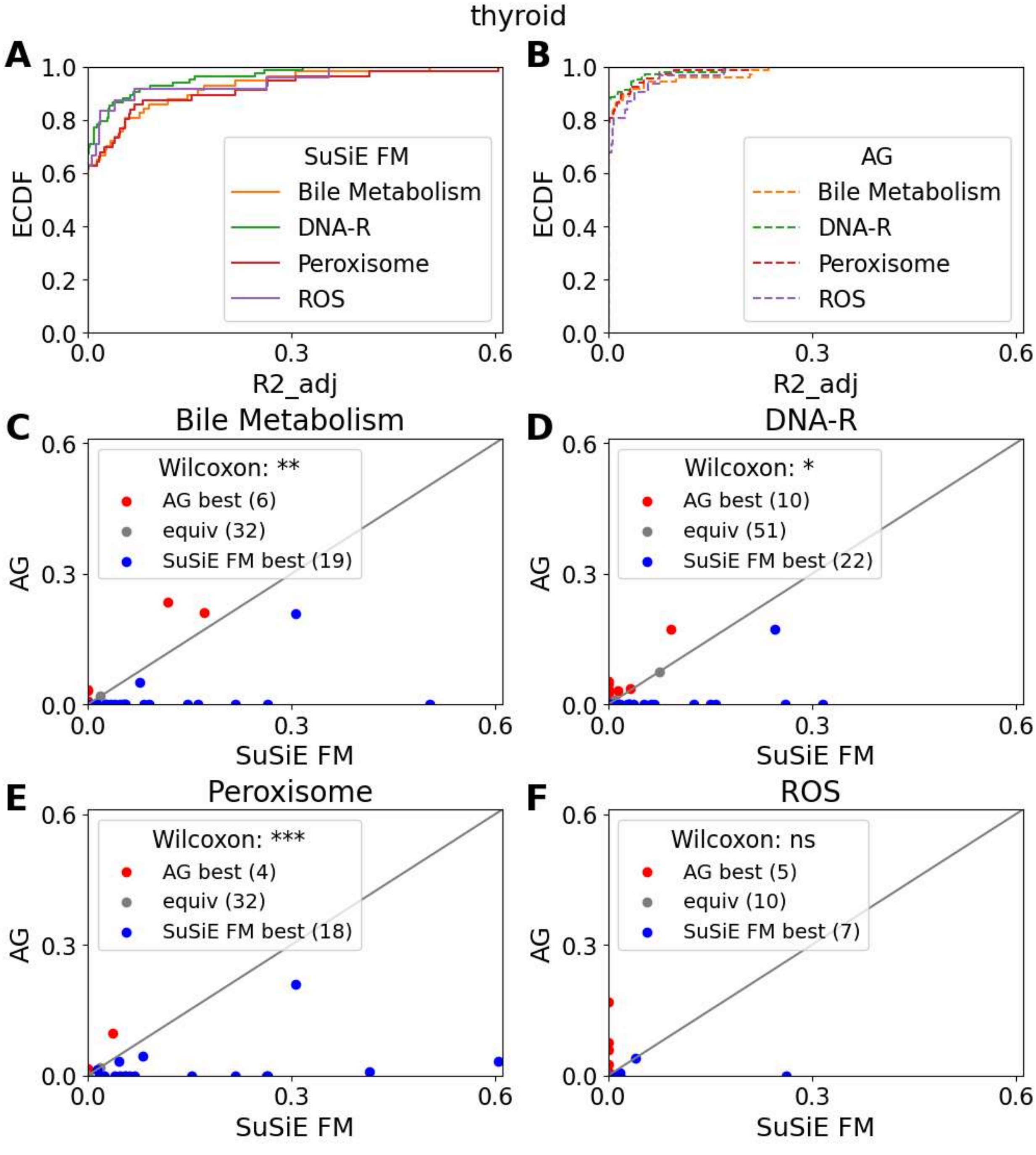


EAGLES coverage in MSigDB Hallmark gene sets. ECDF curves show EAGLES FM (A) and AG (B) performance (adjusted $R^2$) in the independent thyroid cohort over the top 4 gene sets by fraction of members included in EAGLES. For genes modelable in both EAGLES FM and EAGLES AG, scatter plots compare performance in bile metabolism (C), DNA repair (D), peroxisome (E), and reactive oxygen species (F) genes. The Wilcoxon signed-rank test was used for comparisons: ***<0.001; **<0.01; *<0.05; ns>0.05. FM=finemapped; AG=AlphaGenome; DNA-R=DNA repair; ROS=reactive oxygen species.

Figure S27

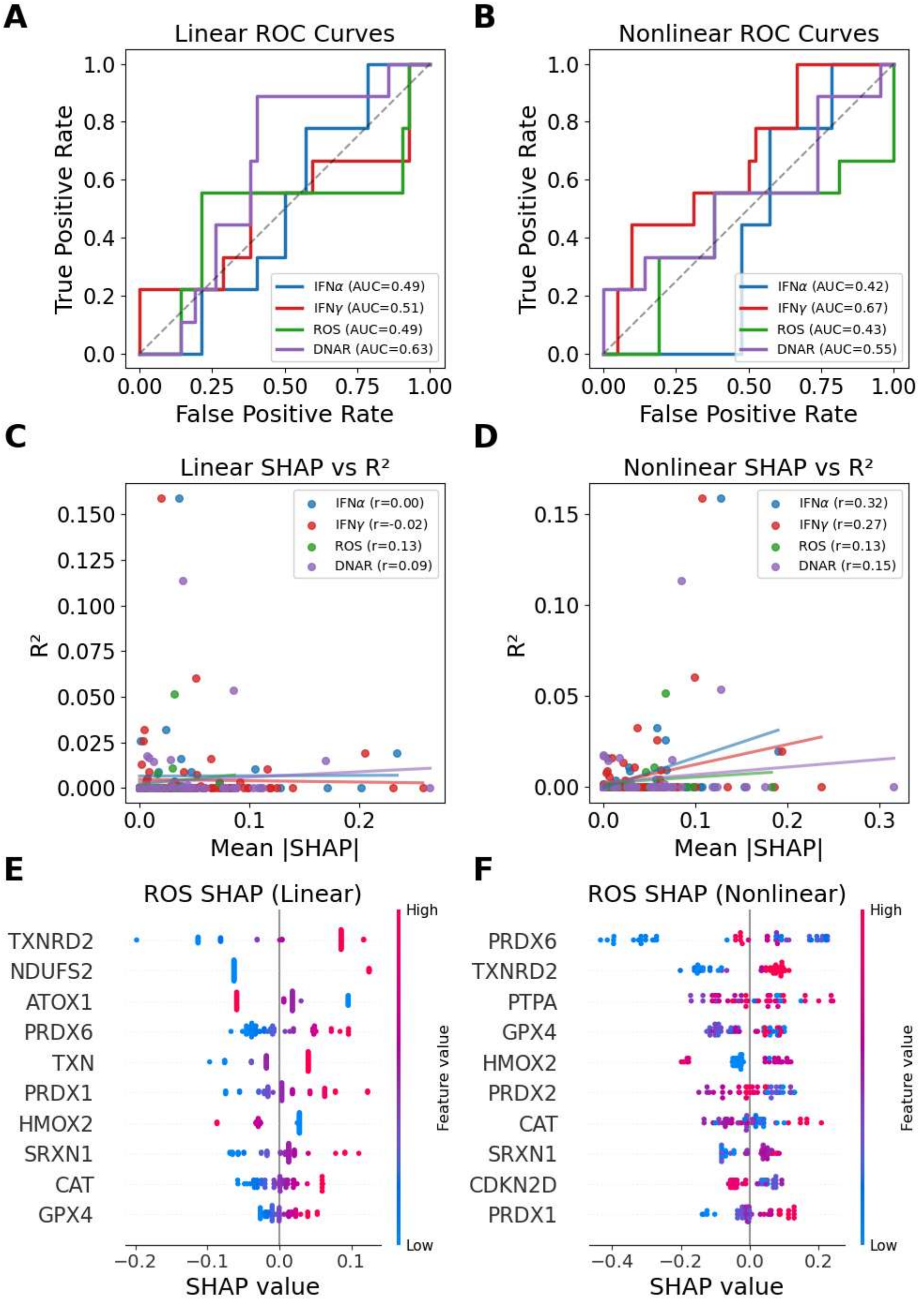


Model performance and feature importance for ICB response prediction using AlphaGenome linear and nonlinear models. ROC curves and AUC values for logistic classifiers trained on IFNα, IFN$\gamma$, ROS, and DNAR gene-score features (A) and XGBoost classifiers trained on the same feature sets (B). Scatter plots show relationships between mean absolute SHAP value and EAGLES adjusted $R^2$ for gene features in logistic (C) and XGBoost (D) classifiers. SHAP beeswarm plots summarize feature importances in the logistic ROS classifier (E) and in the XGBoost ROS classifier (F). Positive SHAP values push models towards predicting response.

Figure S28

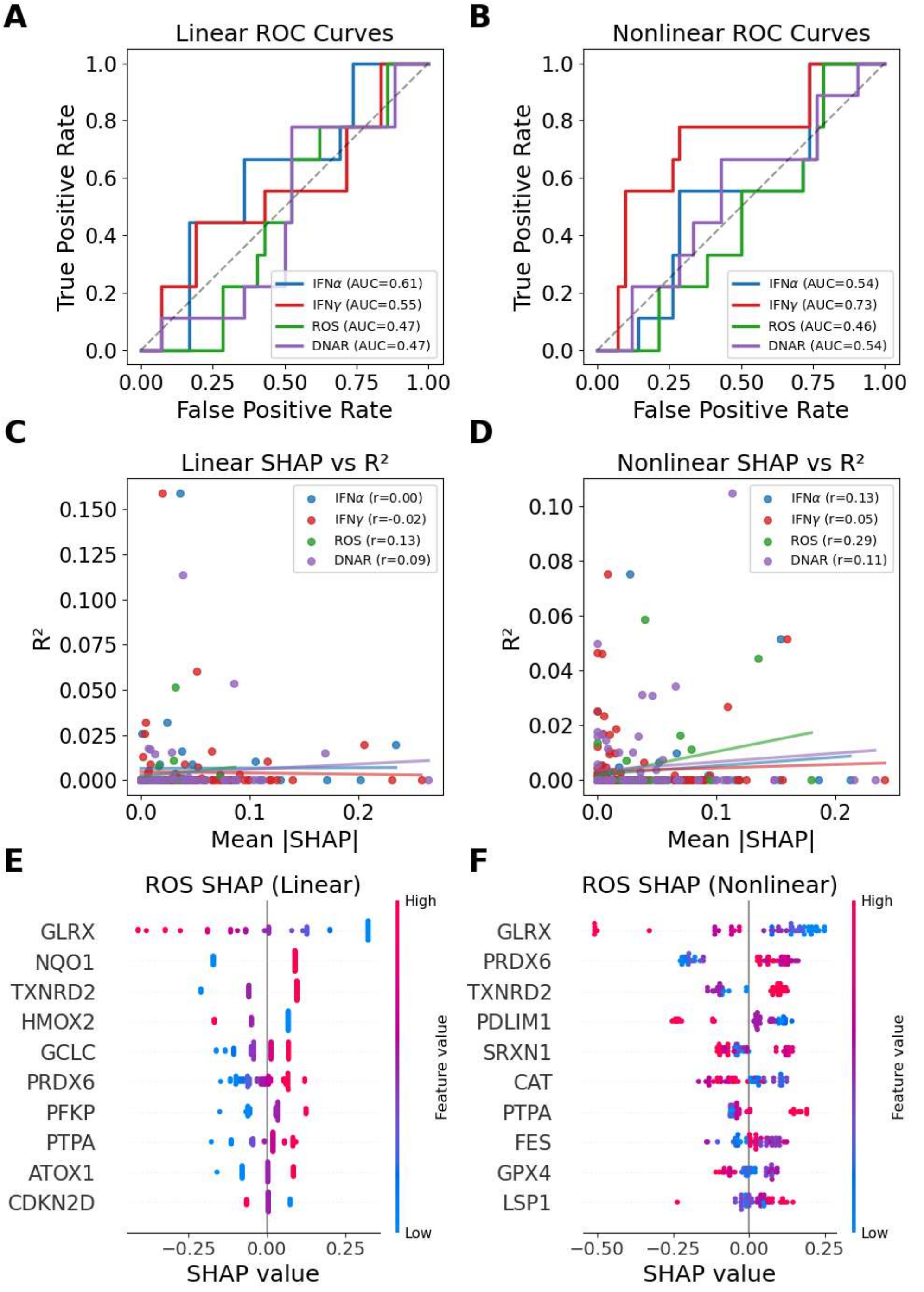


Model performance and feature importance for ICB response prediction using eQTL linear and nonlinear models. ROC curves and AUC values for logistic classifiers trained on IFNα, IFN$\boldsymbol{\gamma}$, ROS, and DNAR gene-score features (A) and XGBoost classifiers trained on the same feature sets (B). Scatter plots show relationships between mean absolute SHAP value and EAGLES adjusted $R^2$ for gene features in logistic (C) and XGBoost (D) classifiers. SHAP beeswarm plots summarize feature importances in the logistic ROS classifier (E) and in the XGBoost ROS classifier (F). Positive SHAP values push models towards predicting response.

Figure S29

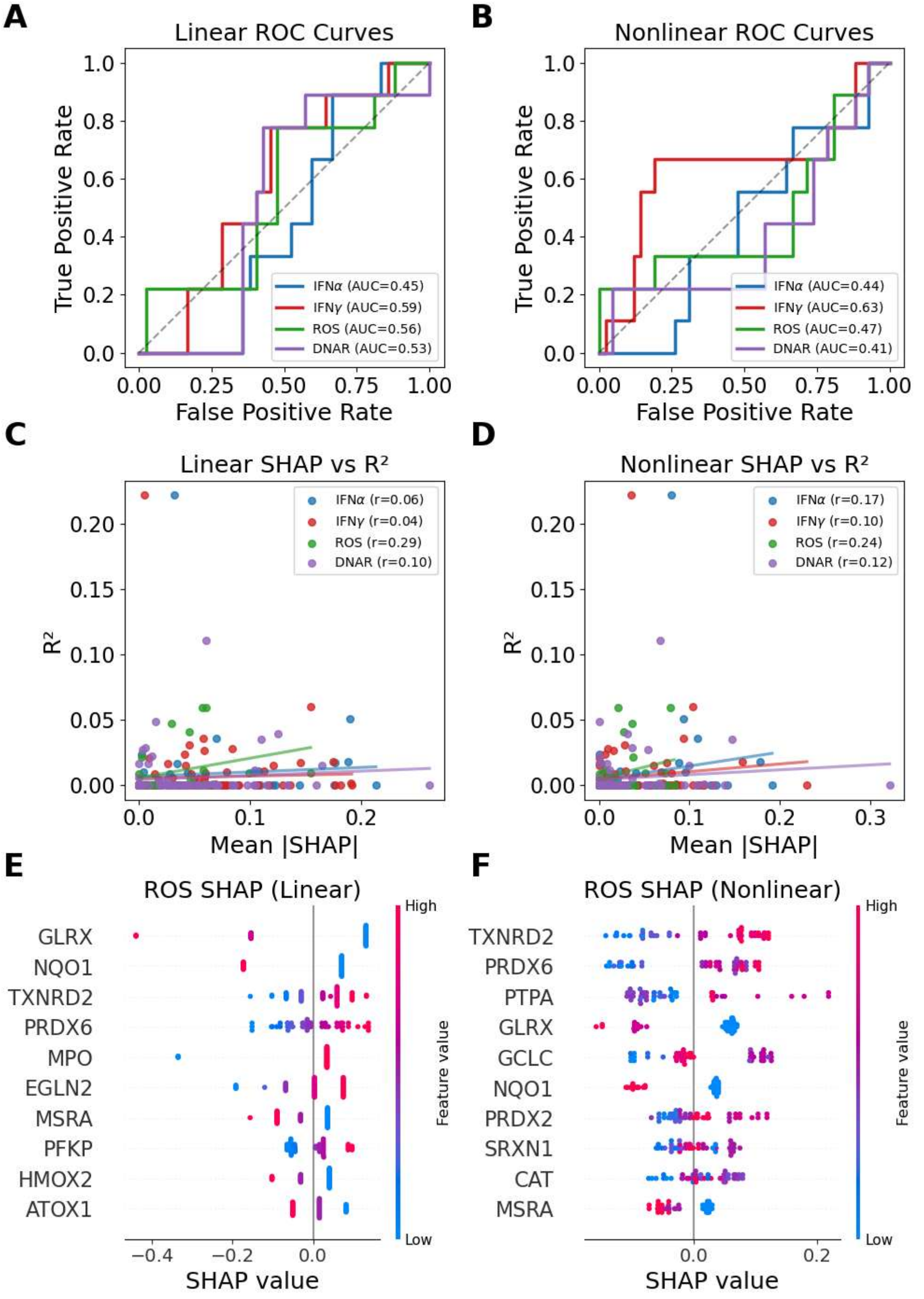


Model performance and feature importance for ICB response prediction using finemapped or AlphaGenome linear and nonlinear models. ROC curves and AUC values for logistic classifiers trained on IFNα, IFN$\gamma$, ROS, and DNAR gene-score features (A) and XGBoost classifiers trained on the same feature sets (B). Scatter plots show relationships between mean absolute SHAP value and EAGLES adjusted $R^2$ for gene features in logistic (C) and XGBoost (D) classifiers. SHAP beeswarm plots summarize feature importances in the logistic ROS classifier

(E) and in the XGBoost ROS classifier (F). Positive SHAP values push models towards predicting response.

Figure S30

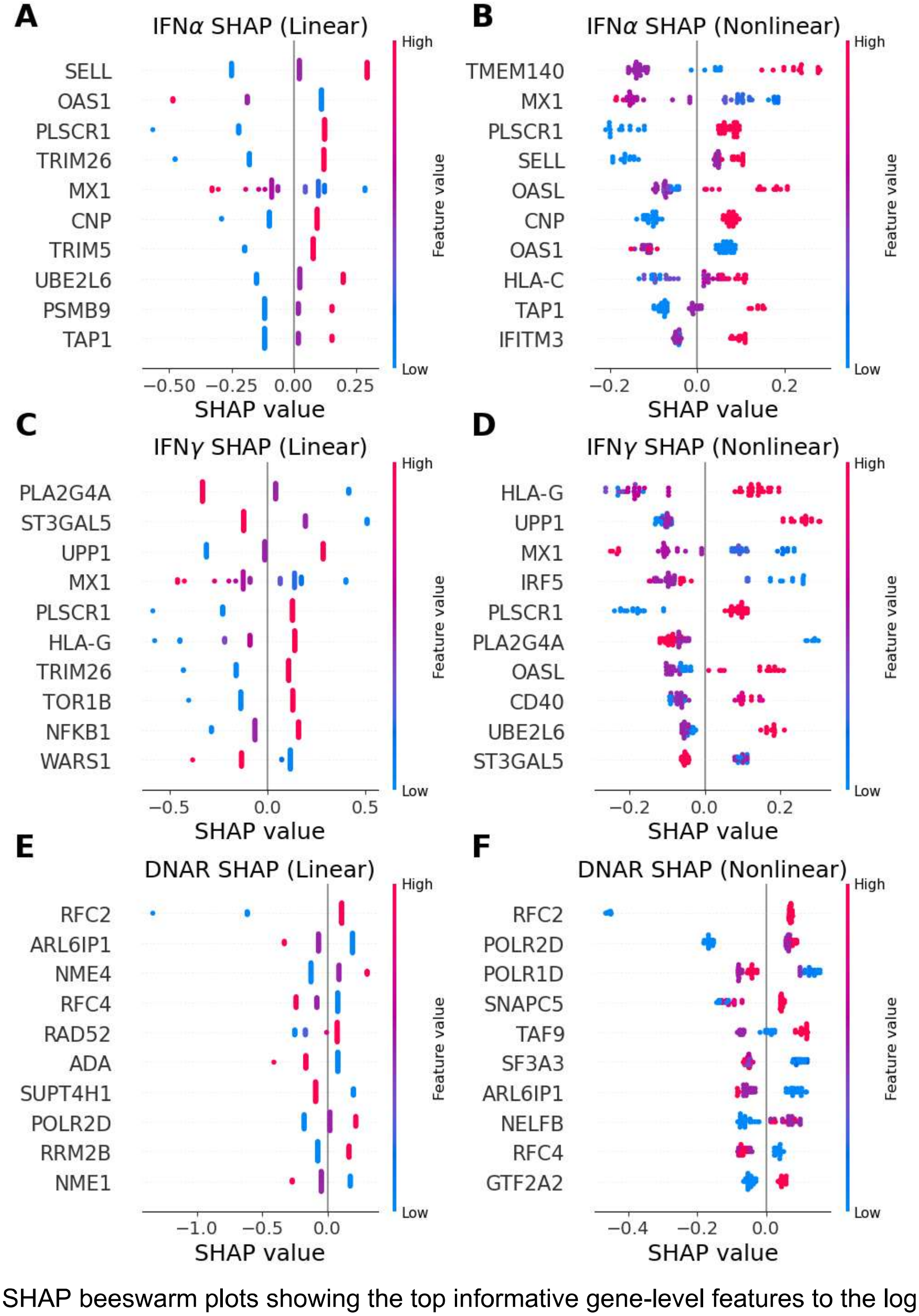


SHAP beeswarm plots showing the top informative gene-level features to the logistic and XGBoost classifiers across the IFNα (A,B) IFN$\gamma$ (C,D), and DNAR (E,F) pathways.
